# Dual-Layer Serological Encoding: Integrating Pathogen-Derived and Host-Reactive Peptide Signatures Reveals Complementary Dimensions of HIV Immune Response

**DOI:** 10.64898/2026.04.20.719572

**Authors:** Dimitry Schmidt, Sergey Biniaminov, Nathalie Biniaminov, Clemens von Bojnicic-Kninski, Roman Popov, Josef Maier, Hubert Bernauer, Johanna Griesbaum, Nicole Schneiderhan-Marra, Alex Dulovic, Alexander Nesterov-Mueller

**Affiliations:** Institute of Microstructure Technology, Karlsruhe Institute for Technology, Hermann-von-Helmholtz-Platz 1, 76344 Eggenstein-Leopoldshafen, Germany; HS Analysis, Haid-und-Neu-Straße 7, 76131 Karlsruhe, Germany; axxelera, Karl-Floesser-Str. 14, 76137 Karlsruhe, Germany; ATG:biosynthetics GmbH, Weberstraße 40, 79249 Merzhausen, Germany; IStLS, Härlestraße 24/1, 78727 Oberndorf a.N., Germany; NMI Natural and Medical Sciences Institute at the University of Tübingen, Markwiesenstraße 55, 72770 Reutlingen, Germany

## Abstract

Serological diagnostics traditionally rely on pathogen-derived antigens to detect infection-specific antibody responses. Chronic infections also induce systemic immune remodeling that may be reflected in global antibody reactivity patterns beyond antigen specificity. Here we evaluate a dual-layer serological framework combining HIV-derived peptides with a host-derived peptide library designed to capture distributed antibody reactivity patterns. Using strict nested cross-validation in a cohort of 105 individuals, pathogen-derived 12-mer peptides achieved high classification performance with an AUC of 0.891, whereas the 10-mer host-based peptide library alone yielded moderate but statistically significant discrimination with an AUC of 0.805. Integration via regularized stacking resulted in only a modest additive improvement, reaching an AUC of 0.897, indicating partial redundancy in diagnostic ranking. In contrast, entropy and inequality analyses revealed substantial immune repertoire restructuring in HIV-positive individuals, characterized by reduced Shannon entropy and significant correlations between classifier probability and repertoire concentration. These findings support a dual-layer model of serology in which the integration of pathogen-derived and host-derived peptides into a meta model encode antigen specificity, whereas host-reactive signatures reflect systemic immune topology. Distinguishing diagnostic ranking from immune-state encoding provides a conceptual framework for multi-layer serological diagnostics.

## 1. Introduction

Serological diagnostics have historically been grounded in antigen specificity, relying on pathogen-derived proteins to detect corresponding antibody responses. In HIV infection, fourth-generation immunoassays measure antibodies directed against viral antigens such as gp41 and p24, achieving high analytical performance through direct epitope recognition (1–3). This antigen-centric paradigm assumes that the dominant serological information is encoded via pathogen-specific antibody binding.

However, chronic viral infections induce systemic immune remodeling that extends beyond antigen specificity (4). HIV infection is characterized by sustained immune activation, B-cell hyperactivation, clonal expansion, altered germinal center dynamics, hypergammaglobulinemia, and perturbations in antibody repertoire diversity (5–9). These systemic alterations suggest that infection may be encoded not only through recognition of viral epitopes but also through structural changes in the global antibody binding landscape.

Peptide libraries designed to resemble host proteomic features provide a complementary strategy to interrogate this systemic immune dimension. Rather than presenting predefined viral antigens, host-related peptide ensembles probe the geometry of antibody binding across a broad sequence space. The resemblance-ranking peptide library concept demonstrated that peptides selected based on proteome-mimetic criteria can enable peptidomic-scale characterization of antibody interaction patterns (10). In that study, high-density peptide arrays were used to identify binding determinants of the therapeutic monoclonal antibody rituximab through ranking-based analysis and motif enrichment, highlighting that antibody interaction structure can be extracted from large peptide landscapes without direct epitope matching (10).

More broadly, immunosignature and high-density peptide array approaches have shown that antibody binding distributions across diverse peptide libraries can discriminate disease states, implying that systemic immune topology — rather than single-epitope recognition — contains diagnostic information (11–14). These findings suggest that host-related peptide landscapes may encode aspects of immune repertoire organization, including inequality, clonal dominance, and diversity collapse. Specifically, while pathogen-specific peptides measure targeted epitope recognition (15, 16), host-related peptide ensembles may capture immune topology — including repertoire concentration and inequality. It remains unclear whether these layers primarily provide redundant diagnostic ranking or whether they encode partially independent biological axes of immune response. To address this, we implement a strict nested cross-validation framework integrating pathogen-derived HIV peptides with a host-related high-density peptide library (**Fig. 1**).

**Figure 1:**
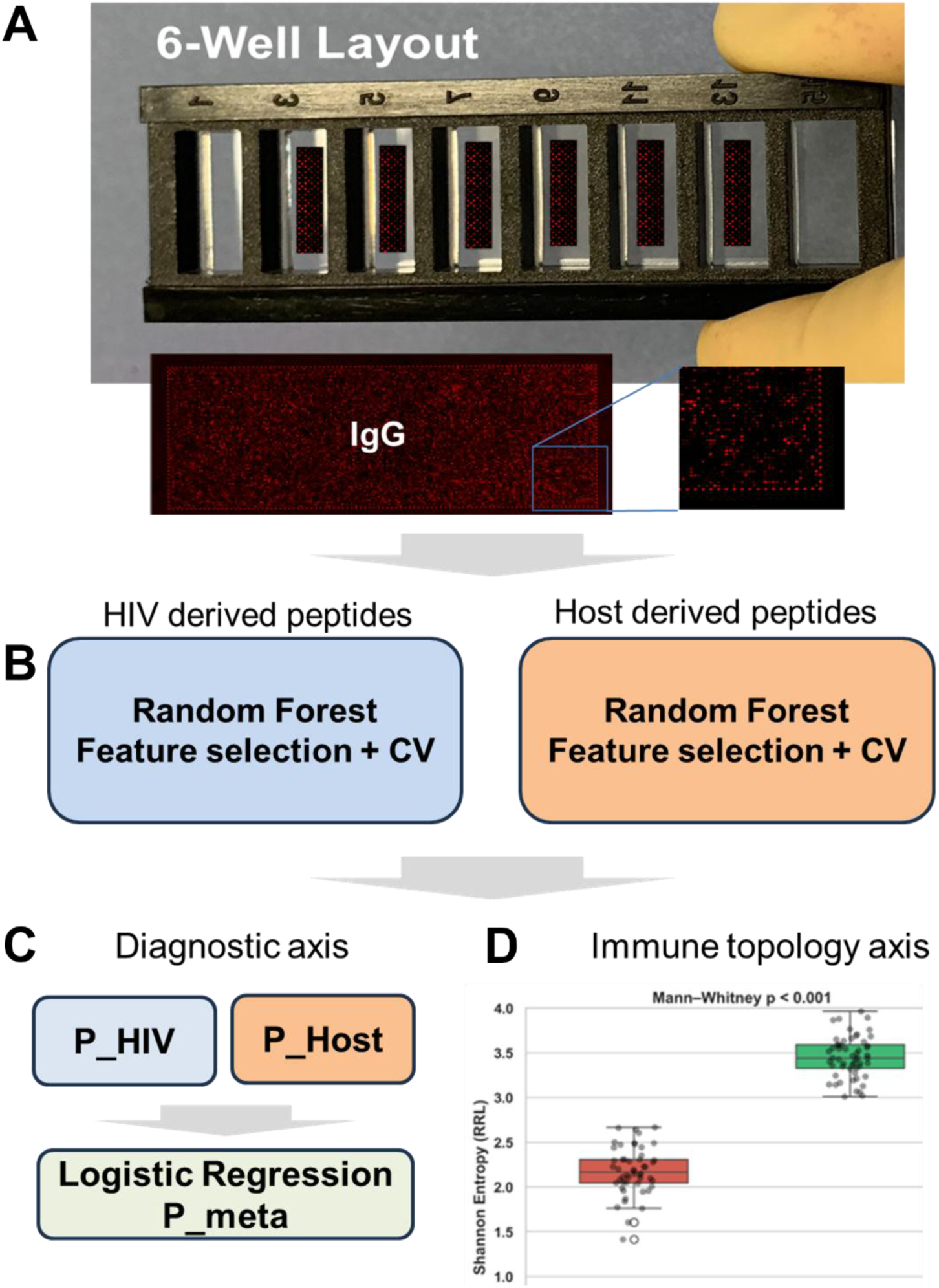
Dual-layer serological framework for HIV classification using pathogen-derived and host-derived peptide signatures. (A) Peptide microarray chip in a 6-well slide format. Each well contains a high-density peptide array used to analyze a single serum sample; the enlarged inset shows spatially resolved IgG fluorescence signals acquired following serum incubation and secondary anti-human IgG labelling. (B) Independent Random Forest classifiers were trained on HIV-derived peptide features (left; 49 peptides, blue) and host-derived resemblance-ranking library (RRL) peptide features (right; 2,087 peptides, orange), each with nested cross-validation and embedded feature selection. (C) Out-of-fold predicted probabilities from both classifiers (P_HIV and P_Host) were combined as inputs to a regularized logistic regression meta-model (P_meta), integrating both serological layers at the probability level. (D) Distributional diversity metrics, including Shannon entropy, were derived from the host peptide library fluorescence distribution to characterize the structural organization of the antibody binding profile independently of diagnostic classification.

By separating diagnostic classification performance from immune topology metrics such as Shannon entropy and Gini index, we evaluate whether multi-layer serology reveals distinct and partially orthogonal dimensions of immunological encoding.

## 2. Methods

### 2.1 Study Cohort

A total of 105 serum samples were analyzed. Serum samples were obtained from the FIND Foundation Biobank, which provided the corresponding HIV status and clinical annotations. The cohort consisted of 35 HIV-positive (HIV+) and 70 HIV-negative (HIV−) individuals (ratio 1:2), giving a balanced two-class structure suitable for stratified cross-validation. Samples were collected from three geographically distinct cohorts: South Africa (n = 37), Peru (n = 37), and Viet Nam (n = 31). Geographic diversity was retained to assess generalizability across HIV-1 subtype distributions and epidemiological settings. Only HIV status was used as the classification endpoint. All samples were stored and processed according to standardized biobank procedures prior to peptide array analysis.

### 2.2 Peptide Libraries

Two peptide libraries were investigated, each targeting a distinct dimension of serum antibody reactivity.

The **pathogen-derived panel** consisted of 49 HIV-derived 12-mer synthetic peptides representing immunologically characterized regions of viral structural proteins: 33 peptides from the gp41 envelope protein (plus 1 randomized control peptide), tiling the major immunodominant region between heptad-repeat sequences and parts of the membrane-proximal external region (MPER), and 13 peptides from the p24 Gag protein (plus 2 randomized controls). Peptides were selected based on published immunodominance data and were designed to capture the dominant IgG responses elicited during natural HIV-1 infection (see Supplementary Methods S1).

The **host-related resemblance-ranking library (RRL)** consisted of 2,087 synthetic 10-mer peptides selected on the basis of compositional similarity to human proteomic fragments, following the resemblance-ranking design principle described by Jenne et al. (10).

### 2.3 Peptide Array Fabrication and Signal Acquisition

Peptide libraries were synthesized in situ on standard glass microscope slides (25 mm × 75 mm) by Axxelera (Karlsruhe, Germany).

Peptide arrays were assembled into 6-window chambers (**Fig.1A**) and hydrated with 500 µL PBS-T (v/v 0.05% Tween20) and incubated for 10 min on a shaker. The buffer was then removed, with the peptide arrays then washed briefly 3x with 500 µL PBS-T. Serum samples were diluted 1:1000 in assay buffer (PBS-T supplemented with 10% (v/v) Rockland Blocking Buffer MB-070), with 400 µL of diluted sample then added to each window. Arrays were then incubated in darkness for 2 hours at room temperature on a shaking incubator (120 mot/min). To remove unbound antibodies, peptide arrays were initially washed quickly 4x inside a sterile workbench with 500 µL PBS-T. To ensure thorough washing, 500 µL PBS-T was then added to each chamber, incubated for 10 min on the shaker (180 mot/min) and then removed. This incubation washing step was then repeated once for a total of 6 washing steps. To detect bound antibodies, 400 µL of 1 µg/mL (diluted in assay buffer) goat anti-human IgG (Biozol #109-605-008) was added to each window and incubated in darkness on a shaker for 45 min (120 mot/min). Following this, peptide arrays were again washed as above to remove unbound secondary antibody. Peptide arrays were then washed briefly in 500 µL of ddH2O (10 min incubation on shaker), soft shaken in a Falcon for 5 seconds to remove large droplets, and then carefully dried with an air pistol. Once dried, peptide arrays were placed in light-shielded falcon tubes filled with argon and stored at 4°C until imaging.

Stained peptide arrays were scanned using an Innoscan 1100 AL confocal fluorescence scanner (Innopsys, Carbonne, France) at a resolution of 2 μm, with a PMT gain of 4 and an excitation wavelength of 635 nm. Fluorescence intensities of the peptide spots were quantified using the MAPIX software (Innopsys, Carbonne, France). MAPIX calculates the median fluorescence intensity within each spot area and assigns it to the corresponding peptide. For peptides with multiple replicate spots, the median intensity across replicates (MFI) was computed and used for subsequent analysis. Replicate spots were randomly distributed on the chip to minimize potential local effects on fluorescence signals. These spot intensities were then used for downstream data analysis.

### 2.4 Data Preprocessing

For each sample, peptide MFI values were assembled into a feature matrix of dimensions 105 × 49 (pathogen-derived panel) and 105 × 2,087 (RRL panel). Signal values were converted to numeric arrays and subjected to z-score standardization. Standardization parameters (mean and standard deviation per peptide) were estimated exclusively on training-fold data and applied to the corresponding held-out validation fold. This fold-specific normalization ensures that no information from the test fold can influence feature scaling, preventing a common source of data leakage in high-dimensional cross-validation.

All preprocessing steps, including missing-value handling, feature standardization, and feature selection were embedded within the cross-validation loop and never applied globally across the full dataset.

### 2.5 Machine Learning and Feature Selection

All modeling analyses were performed in Python using the scikit-learn library (version 1.2). To obtain unbiased performance estimates and prevent overfitting in the high-dimensional setting, nested stratified cross-validation was implemented with a 5-fold outer loop and a 5-fold inner loop. The outer loop was used exclusively for performance estimation on held-out data, while the inner loop was used for feature selection and model hyperparameter optimization.

Within each outer training fold, feature ranking was performed using Random Forest classifiers with class-balanced sample weighting (class_weight=’balanced’), applied to compensate for the 1:2 HIV+/HIV− imbalance. Feature importance was determined using Gini impurity-based importance scores aggregated across all trees in each forest.

Candidate feature subset sizes *k* were evaluated systematically within the inner cross-validation loop. For each candidate *k*, the top-ranked features were selected and classification performance was assessed using the mean inner-fold AUC. Adaptive *k*-selection was performed by identifying the smallest *k* whose mean inner-fold AUC was within a 1% tolerance of the best-observed AUC.

This parsimonious selection criterion favors compact feature panels, reducing the risk of incorporating redundant or noise-amplifying features while maintaining near-maximal predictive performance.

The optimized feature subset and trained classifier were then evaluated on the held-out outer test fold to generate out-of-fold (OOF) predicted class probabilities. Stable features were defined as peptides selected in at least 60% of the five outer cross-validation folds (i.e., in ≥3 of 5 folds). Recursive feature elimination with cross-validation (RFECV) (17, 18) was additionally applied as an independent feature-elimination method to estimate optimal subset size and validate the adaptive *k*-selection results.

### 2.6 Meta-Model Integration

To quantify information overlap and complementarity between the two serological layers, Pearson correlation was computed between OOF predicted probabilities from the pathogen-derived and RRL classifiers. Bootstrap resampling (10,000 iterations, sampling with replacement) was used to derive 95% confidence intervals for the correlation coefficient. Scatter plots and kernel density estimates were generated to characterize probability alignment and distributional structure.

To assess combined diagnostic performance, a regularized logistic regression meta-classifier was trained on the probability outputs of both primary models. The meta-feature matrix *X_meta_* was defined as:

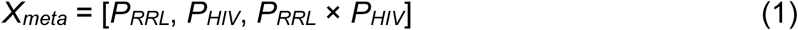

where *P_RRL_* and *P_HIV_* are the OOF predicted probabilities from the RRL and pathogen-derived models respectively, and the third term is a pairwise interaction feature. The ground-truth label was HIV status.

Meta-classification was performed using stratified 5-fold outer cross-validation. Within each outer fold: (i) training data were standardized using StandardScaler (18) fitted on training data only; (ii) a logistic regression classifier (solver = lbfgs, max_iter = 2,000) was trained; (iii) the regularization parameter *C* was optimized using GridSearchCV within a 3-fold inner loop over the grid:

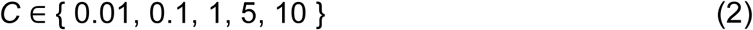

Model selection within the inner loop was based on ROC AUC. The optimized meta-model was then applied to the held-out outer fold to generate meta-level OOF probabilities. At no stage did the meta-model have access to in-fold primary model predictions, ensuring no data leakage between the primary and meta-classification stages.

Final performance was evaluated using pooled OOF predictions. The integrated gain was defined as:

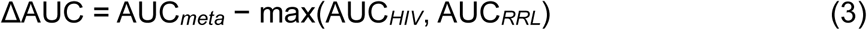

### 2.7 Immune Distribution Metrics and Statistical Analysis

To characterize structural properties of antibody binding within the host-related peptide layer, five per-sample distributional metrics were computed across all 2,087 RRL peptide signals. For each sample, raw fluorescence intensities *x_i_* (for peptide *i* = 1,…, *p*, where *p* = 2,087) were converted to relative proportions:

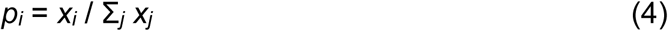

This normalization ensures that each sample’s signal distribution sums to unity, enabling comparison of distributional shape independently of absolute signal magnitude.

Shannon entropy was computed to quantify the diversity of the peptide binding distribution, with higher values indicating more uniform spread across peptides and lower values indicating concentration of binding in fewer peptides:

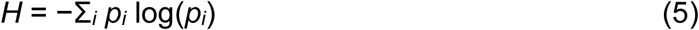

Natural logarithms were used throughout.

The Gini index was computed to quantify distributional inequality, with higher values indicating stronger concentration of signal in a subset of peptides:

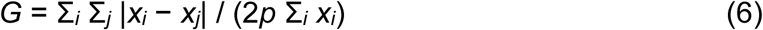

*G* ranges from 0 (perfect equality across all peptides) to 1 (maximum inequality, all signal concentrated in one peptide).

Signal variance was computed as the mean squared deviation of peptide intensities from the sample mean:

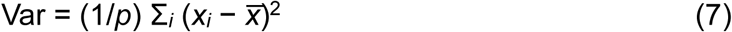

where *x̅* denotes the within-sample mean fluorescence intensity.

Top-peptide dominance was defined as the proportion of total signal contributed by the single highest-intensity peptide:

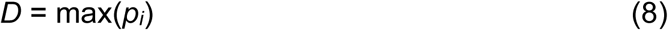

This metric captures extreme distributional skew attributable to a single dominant binding partner.

Between-group comparisons of all five distributional metrics (*H*, *G*, Var, *D*, and mean signal) between HIV+ and HIV− samples were performed using the two-sided Mann–Whitney *U* test, which makes no assumptions about distributional normality. Effect sizes were quantified using Cliff’s δ, interpreted as small (δ = 0.147), medium (δ = 0.33), and large (δ = 0.474) following published thresholds (19).

Associations between RRL classifier predicted probabilities and distributional metrics were evaluated using both Pearson (linear) and Spearman (monotonic) rank correlation coefficients. Bootstrap resampling (10,000 iterations) was applied to estimate 95% confidence intervals for all correlation coefficients. All statistical tests were two-sided with α = 0.05. All analyses were implemented in Python using NumPy, SciPy, and scikit-learn.

## 3. Results

### 3.1 Pathogen-Derived Peptides Dominate Diagnostic Classification

Nested 5×5 cross-validation demonstrated strong diagnostic performance of the pathogen-derived peptide panel (49 peptides). The pooled out-of-fold area under the ROC curve (AUC) was 0.891 (**Fig. 2A**), with sensitivity 0.83 and specificity 0.94 at the Youden-optimal threshold.

**Figure 2:**
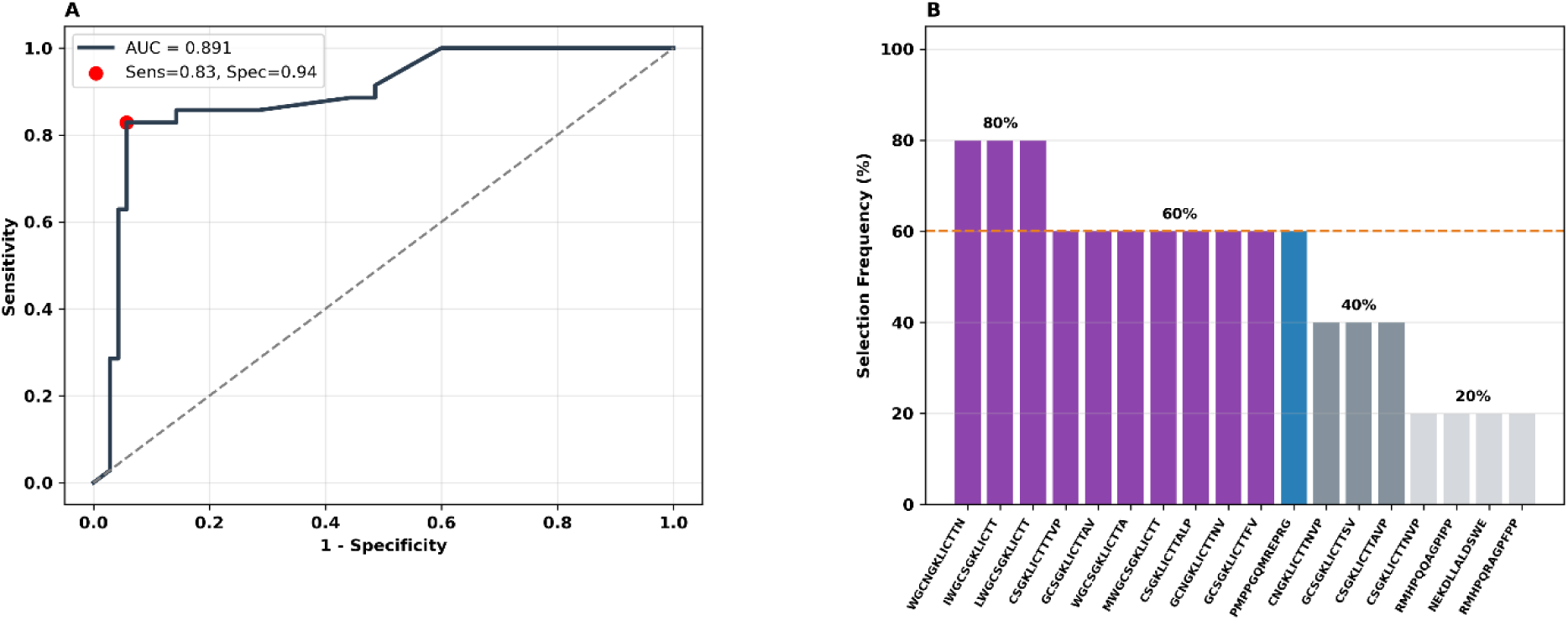
HIV-derived peptide panel identifies a stable gp41 HR-linker serological signature. (A) Receiver operating characteristic (ROC) curve for the Random Forest model under nested 5-fold leave-one-out cross-validation (n = 105; HIV+ = 35, HIV− = 70). The operating point (filled circle) was selected at the Youden-optimal threshold (0.611). AUC = 0.891; sensitivity = 83%; specificity = 94%. (B) Peptide stability frequency plot showing the proportion of outer cross-validation folds (out of 5) in which each of the 49 screened peptides was included in the optimal panel. The dashed line indicates the 60% stability threshold. Purple bars = gp41 HR1-HR2-linker derived stable peptides (n = 10); blue bar = p24 Gag stable peptide (n = 1); grey bars = frequently selected (40–59%); light grey = infrequent (<40%).

Feature reduction consistently selected ten peptides are from gp41 major immunodominant region epitope (588-607 in Env of HBX2, all covering the region 595-609; the region corresponds to the 46-residue highly flexible linker sequence between the conserved three-helix bundles of the heptad-repeat regions HR1 and HR2) and one peptide from p24 (90-101; this region is in the N-terminal domain, in the cyclophilin A-binding region), meeting stability criteria (≥60% outer-fold selection frequency) (**Fig. 2B**). Recursive feature elimination indicated convergence toward a compact performance plateau (Supplementary Fig. S1), with statistical significance confirmed by permutation testing (p = 0.0099). Despite maximal performance at higher feature counts, a 15-peptide panel achieved equivalent diagnostic accuracy within 1% of the maximal AUC (Supplementary Fig. S2). Considering this plateau and the biological redundancy within the gp41 heptad-repeats (HR1, HR2) linker region, the parsimonious 15-peptide panel was selected for downstream analyses.

To visualize signal structure within the pathogen-derived peptide panel, we constructed a heatmap of the 11 stable peptides selected in ≥60% of outer cross-validation folds (**Fig. 3**). The heatmap reveals a highly coherent binding pattern across HIV-positive individuals. Ten of the eleven stable peptides derive from gp41 (env), forming a tightly clustered signal block characterized by strong and consistent enrichment in HIV-positive samples. In contrast, HIV-negative samples exhibit uniformly low z-scored signal intensities across these peptides, indicating high specificity.

**Figure 3:**
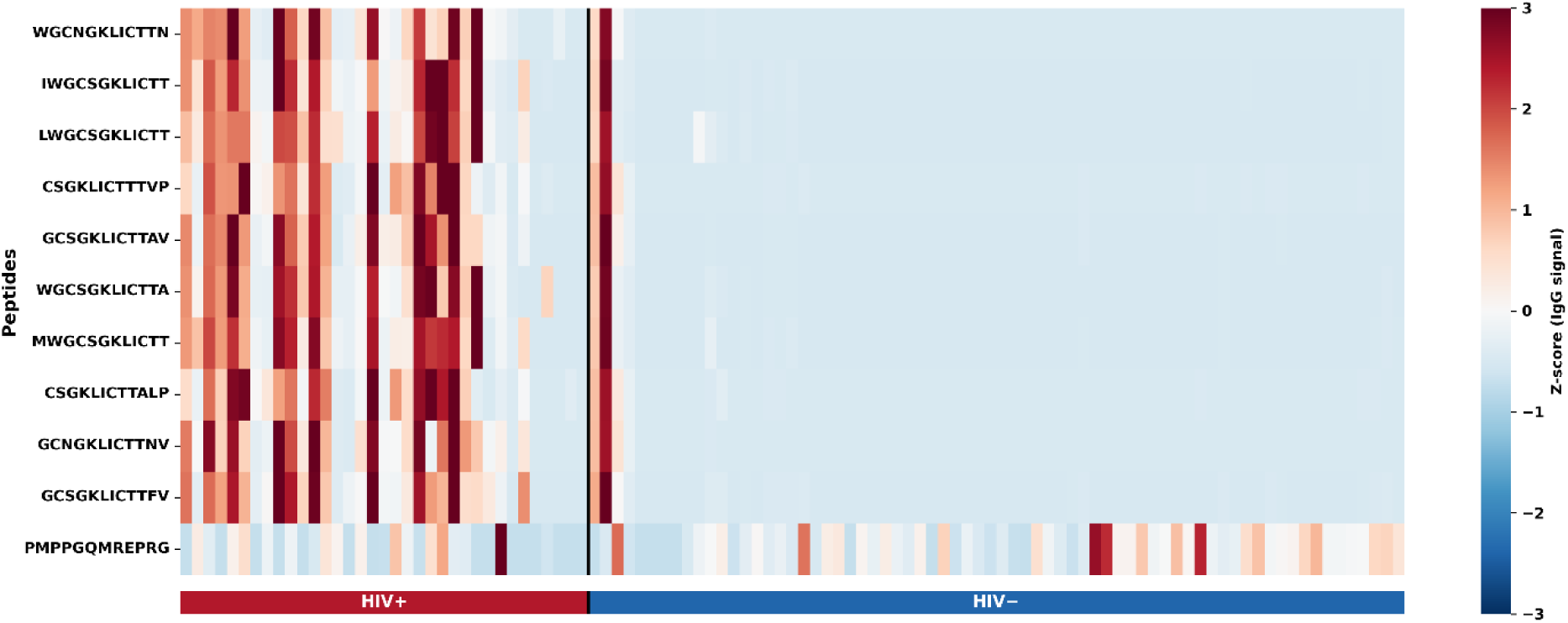
Z-scored IgG reactivity across the stable peptide panel. Heatmap of Z-scored IgG signal intensities for the 11 stable peptides selected by nested cross-validation (≥3/5 outer folds). Signals were standardized per peptide across the full cohort (n = 105) and clipped at ±3 standard deviations. Samples are ordered by HIV status (HIV+, n = 35; HIV−, n = 70) and sorted within each group by descending predicted probability. The vertical black line separates HIV+ and HIV− samples.

Notably, the gp41-derived peptides share overlapping sequence motifs centered on the structurally constrained immunodominant epitope cluster of the linker region between HR1 and HR2. Signal intensity patterns across these peptides are highly concordant, consistent with recognition of a shared conformational or linear epitope region.

The single p24-derived peptide (PMPPGQMREPRG) displays a distinct signal distribution compared to the gp41 cluster, showing lower magnitude and greater heterogeneity. This indicates that while p24 contributes to classification, the dominant diagnostic signal is encoded in gp41.

In contrast, the host-related RRL peptide layer (2,087 peptides) achieved moderate classification performance (AUC = 0.805; **Fig. 4A&B**; Supplementary Fig. S3), indicating partial but weaker diagnostic encoding compared to pathogen-specific peptides.

**Figure 4:**
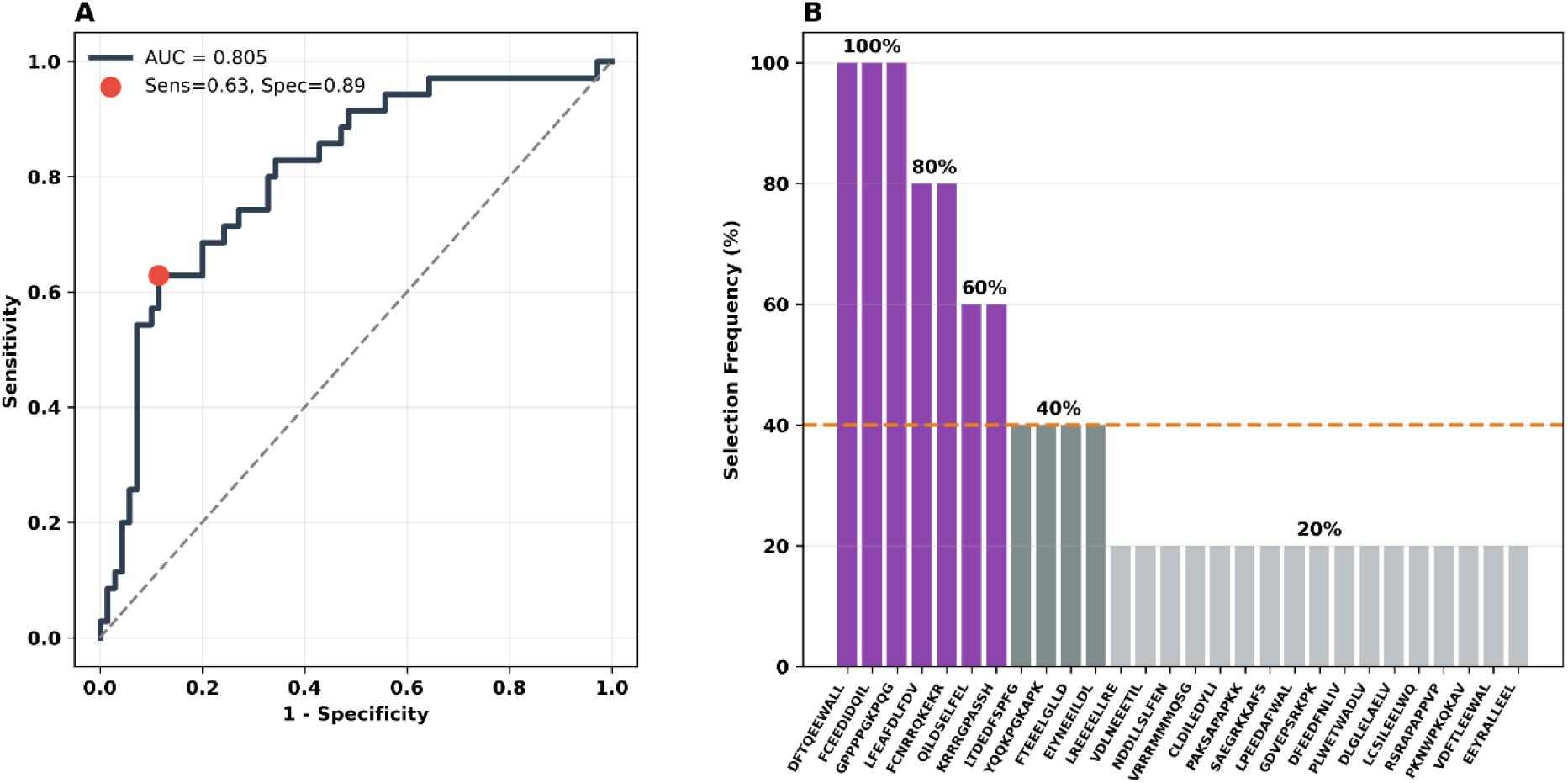
Host-related peptide layer captures stable serological motifs associated with HIV infection. (A) Receiver operating characteristic (ROC) curve for the Random Forest classifier trained on the host-related RRL peptide library under nested 5-fold cross-validation (n = 105; HIV+ = 35, HIV− = 70). The operating point (filled circle) corresponds to the Youden-optimal threshold. AUC = 0.805; sensitivity = 63%; specificity = 89%. (B) Peptide stability frequency plot showing the proportion of outer cross-validation folds (out of 5) in which each peptide was included in the optimal feature subset. The dashed line marks the 40% inclusion threshold. Three peptides were selected in 100% of folds, indicating maximal stability across cross-validation splits. These top-ranked sequences are enriched in proline-containing motifs and low-complexity structural elements, suggesting recognition of conformationally constrained or polyproline-like epitopes. Additional peptides selected in ≥60% of folds include mixed hydrophobic and charged motifs, consistent with recognition of structural antibody-binding geometries rather than canonical linear pathogen epitopes. Dark purple bars indicate highly stable peptides (≥60%); dark grey bars represent moderately stable features (40–59%); light grey bars correspond to infrequently selected peptides (<40%).

### 3.2 Probability-Level Integration Yields Modest Gain

To assess cross-layer complementarity, out-of-fold probabilities from both classifiers were integrated using a regularized logistic regression meta-model (see Methods). The integrated model achieved an AUC of 0.897, corresponding to a marginal improvement over the pathogen-derived panel (ΔAUC = +0.006). ROC comparison (Supplementary Fig. S4) demonstrated near-overlapping performance profiles, indicating that integration provides only limited additional discriminatory information.

### 3.3 Partial Orthogonality Between Serological Layers

To assess cross-layer overlap, Pearson correlation was calculated between out-of-fold predicted probabilities from the pathogen-derived and host-related classifiers. A moderate correlation was detected (r = 0.55; 95% bootstrap CI: 0.387–0.697), corresponding to ∼30% shared variance (r² ≈ 0.30) (**Fig. 5A**). The observed dispersion around the identity line indicates that a substantial fraction of ranking information is not shared between layers. Density analyses further demonstrate distinct probability distributions (**Fig. 5B**), indicating differences in score structure beyond linear association. Collectively, these results support partial orthogonality rather than redundancy, suggesting that the two layers capture complementary aspects of the serological response.

**Figure 5:**
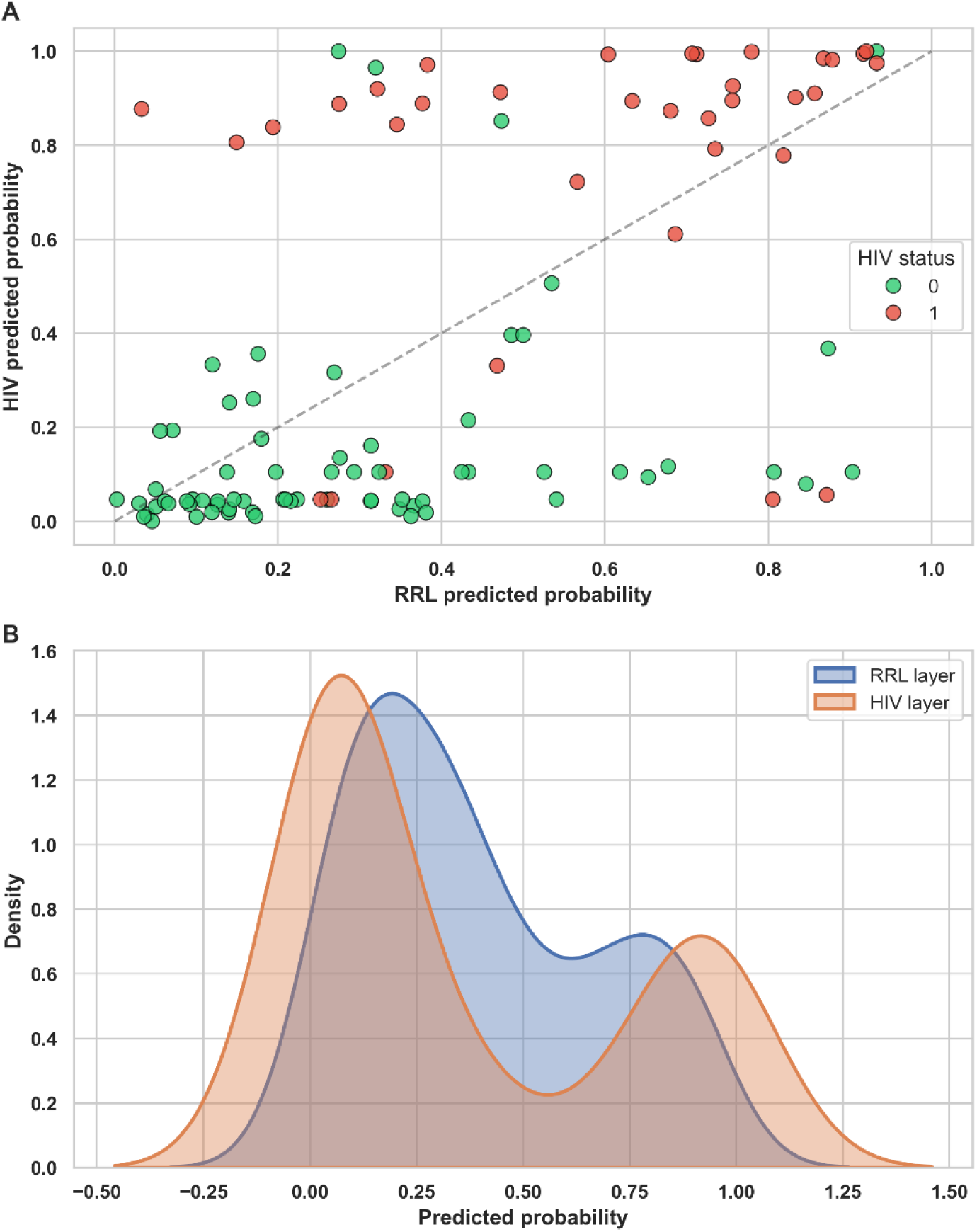
Partial orthogonality of serological classification layers. (A) Correlation of out-of-fold predicted probabilities from the RRL and pathogen-derived classifiers (n = 105). A moderate correlation was observed (r = 0.55; 95% CI: 0.387–0.697; r² ≈ 0.30), indicating incomplete overlap in diagnostic ranking. (B) Density distributions of classifier probabilities reveal distinct score structures between layers. These findings demonstrate partial orthogonality, consistent with complementary encoding of serological information.

### 3.4 Host-Related Layer Encodes Immune Repertoire Remodeling

To determine whether the host-related layer reflects structural properties of antibody binding rather than purely diagnostic ranking, distributional metrics were computed per sample across the RRL peptide set.

Shannon entropy of the RRL signal distribution was significantly lower in HIV+ compared to HIV− individuals (Mann–Whitney p = 0.012; Cliff’s δ = −0.30), indicating reduced binding diversity (**Fig. 6A**). This pattern is consistent with contraction or concentration of the antibody binding landscape during chronic infection (7).

**Figure 6:**
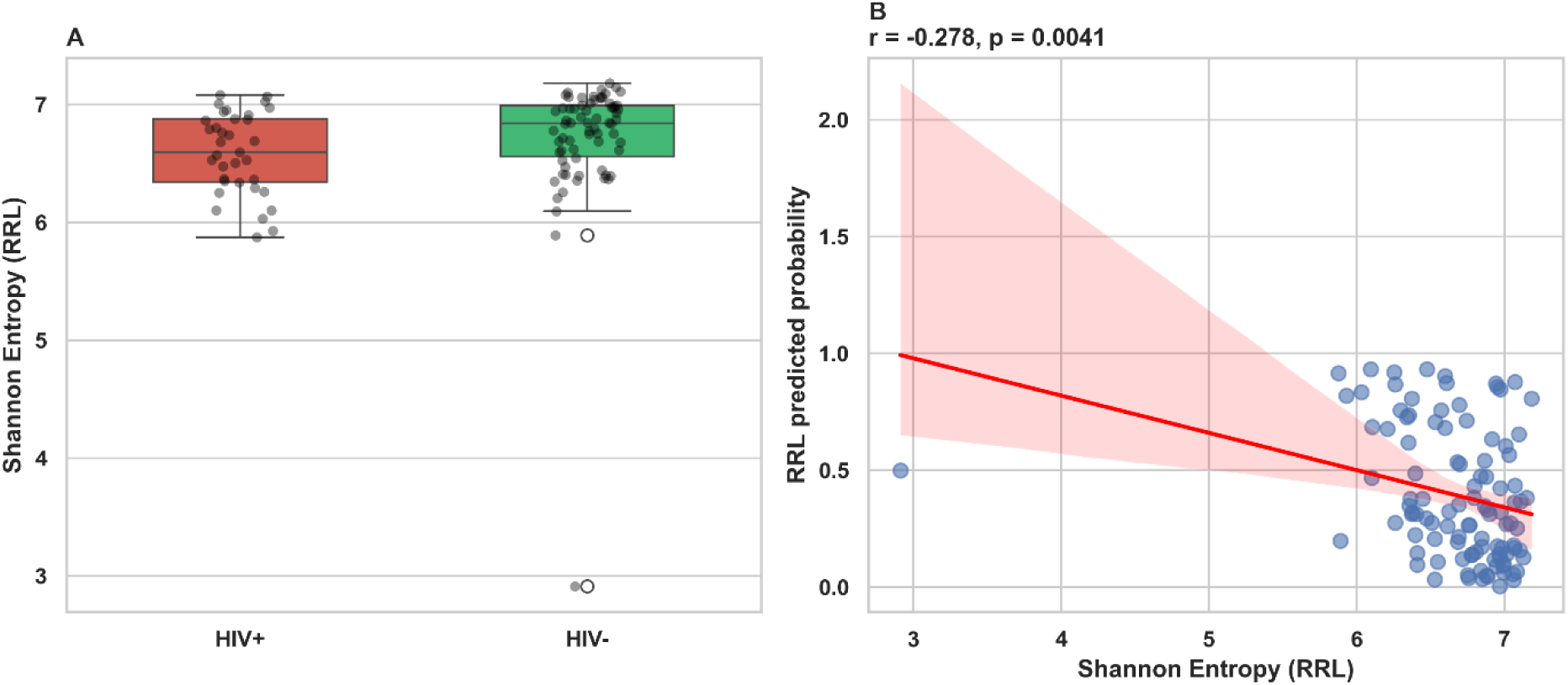
Reduced repertoire entropy and its association with RRL classifier probability. (A) Comparison of Shannon entropy between HIV+ and HIV− individuals (n = 105). HIV+ samples displayed lower entropy (Mann–Whitney p = 0.012; Cliff’s δ = −0.30). Boxes indicate IQR, centre lines show medians, whiskers denote 1.5 × IQR, and individual points represent samples; points beyond whiskers are shown as outliers. (B) Scatter plot of Shannon entropy versus RRL-predicted probability. A significant inverse correlation was observed (Pearson r = −0.278, p = 0.004).

We next evaluated whether these structural metrics were associated with RRL classifier probability. Entropy showed a significant negative correlation with RRL probability (Pearson r = −0.278, p = 0.004) (**Fig. 6B**), whereas the Gini index exhibited a positive correlation (r = 0.376, p < 0.001) (Fig. S5). In contrast, signal variance and top-peptide dominance were not significantly associated with classifier output (Supplementary Fig. S6).

Higher RRL-predicted HIV probability was therefore associated with lower repertoire entropy and greater signal inequality, without corresponding changes in overall signal variance or single-peptide dominance.

## 4. Discussion

### 4.1 Convergence on the gp41 MPER: Data-Driven Rediscovery of an Immunodominant Epitope Cluster

The pathogen-derived classifier converged on a structurally coherent cluster of gp41-derived peptides sharing the conserved CSGKLICTT core motif, with ten of eleven stability-selected peptides mapping to the gp41 envelope protein. This region is part of the primary immunodominant region (PID) on gp41 (20). The PID induces multiple non-neutralizing antibodies and can be considered a decoy structure misleading and dominating the immune response. It is exposed on gp41 stumps on post-fusion virions after shedding of gp120 subunits of the envelope protein. The PID itself is an amphipathic 15-amino acid region (596-610, WGCSGKLICTTAVPW consensus), which separates the two heptad-repeat domains (HR1 and HR2) present in the post-fusion conformation of gp41. It has an internal disulfide bond between C598 and C604. It is nevertheless highly conformational flexible. This flexibility allows both, germline and mature antibody structures to bind and causes high immunodominance, with inducing ∼70 % of all antibodies during an acute HIV-1 infection. The fact that the present classifier independently converges on this region through fully data-driven feature selection serves as an internal validation of both the experimental platform and the modeling framework.

The high signal coherence observed across the overlapping PID-derived peptides (**Fig. 2**) is consistent with recognition of a shared linear or conformationally constrained epitope rather than multiple independent binding specificities. The overlapping sequence design of these peptides tiles a contiguous stretch of the PID, meaning that correlated classifier features reflect genuine biological redundancy: multiple peptides capture the same underlying antibody-antigen interaction. This redundancy reduces effective feature dimensionality while simultaneously increasing classification robustness, as any single-peptide signal dropout is compensated by flanking sequences. From a diagnostic engineering perspective, this supports the use of epitope-tiling arrays for immunodominant regions when signal stability across samples is prioritized.

The single p24-derived stable peptide (PMPPGQMREPRG) presents a distinct biological profile compared to the gp41 cluster. The peptide is part of a long antigenic peptide co-crystallized with the murine monoclonal antibody Fab fragment 25.3 (21) in the N-terminal domain of p24, the core pentapeptide PGQMR was found to be not present in the human proteome (22). Its signal distribution showed lower magnitude and greater inter-individual heterogeneity (**Fig. 2**), consistent with known variability in anti-p24 antibody responses as a function of disease stage and antiretroviral treatment history. The peptide’s proline-methionine-proline-proline (PMPP) N-terminal motif confers structural rigidity through a polyproline-II-like backbone geometry (23), which may restrict its recognition to a specific antibody subpopulation. Importantly, this proline-rich structural character creates a conceptual bridge to the host-related peptide layer, as discussed in the following section.

### 4.2 The RRL Stable Peptide Signatures: Proline-Rich Motifs and the Structural Geometry of Antibody Recognition

The three peptides selected across 100% of outer cross-validation folds did not share an obvious dominant sequence motif. One of the peptides selected in 100% of outer folds contained multiple proline residues. Proline is unique among the canonical amino acids in its pyrrolidine side chain, which imposes rigid backbone dihedral angles (φ ≈ −75°, ψ ≈ 150°) and is incompatible with standard alpha-helical and beta-sheet conformations (23). Proline-rich sequences therefore adopt polyproline-II (PPII) helical structures or extended conformations, presenting a constrained and solvent-exposed geometric surface to antibody complementarity-determining regions (CDRs). PPII motifs are abundant in signaling proteins (SH3 domain ligands), extracellular matrix components, and host-pathogen interaction interfaces (24). The consistent selection of such peptides across cross-validation folds suggests that HIV+ sera contain a stable population of antibodies capable of recognizing this structural geometry.

Several immunological mechanisms may account for this observation. First, HIV-1 encodes proline-rich structural elements in multiple proteins, most prominently in Gag (including the PMPP motif of the selected p24 peptide PMPPGQMREPRG discussed above) and in the cytoplasmic tail of gp41. Antibodies induced by these viral proline-rich regions may cross-react with compositionally similar host-mimetic RRL peptides presenting analogous structural geometries, even in the absence of sequence identity. HIV-1 Nef, another viral protein, contains proline-rich motifs that mediate interactions with host SH3 domain-containing signaling proteins through PPII-mediated binding (24), raising the possibility that anti-Nef antibodies may also contribute to RRL reactivity. This structural mimicry mechanism would explain why the RRL layer achieves moderate diagnostic discrimination without targeting canonical viral epitopes. Second, HIV-associated polyclonal B-cell hyperactivation — a well-documented consequence of sustained antigenic stimulation and dysregulated T-helper signaling (7, 25) — generates a population of antibodies with broad, low-affinity binding to structural epitope geometries. A large fraction of class-switched IgG antibodies recovered from HIV-seropositive donors is polyreactive, recognizing multiple structurally diverse antigens through conformationally flexible heavy-chain CDR3 regions (26). Proline-containing peptides, by presenting a restricted and stereotyped backbone geometry, may preferentially capture this expanded polyreactive antibody pool. Polyreactivity in the anti-HIV antibody response has been shown to be common and conserved, affecting both neutralizing and non-neutralizing antibody classes (27). The partial but significant diagnostic power of the RRL classifier (AUC = 0.805) is consistent with this interpretation: the signal is real and reproducible, but less specific than direct epitope recognition.

Third, proline-rich motifs are highly abundant in host proteins involved in signaling (SH3 domain ligands), extracellular matrix components (collagen triple helix, fibronectin PPII regions (23, 24)), and cytoskeletal architecture. Elevated reactivity to such motifs in HIV+ individuals may reflect a component of bystander autoimmune activation, consistent with clinical observations of increased polyreactive and autoreactive antibody titers in chronic HIV infection (28, 29). This interpretation is speculative in the current dataset, but could be directly evaluated in future studies using parallel autoantibody profiling or SH3-domain ligand competition assays.

### 4.3 Partial Orthogonality and the Limits of Diagnostic Integration

The moderate inter-layer correlation observed between out-of-fold predicted probabilities (Pearson r = 0.55; r² ≈ 0.30) indicates that approximately 70% of the variance in classifier output is not shared between layers. This partial orthogonality is biologically interpretable: pathogen-derived peptides encode direct epitope recognition, a binary-like antigen exposure signal that is either present or absent in each individual, whereas the RRL layer encodes a continuous immune topology signal reflecting the structural organization of the global antibody repertoire. These two dimensions are related — HIV infection causes both — but are not equivalent. The former is informative for diagnostic ranking; the latter reflects the immunological state of the host.

The modest additive gain achieved by probability-level stacking (ΔAUC = +0.006) confirms that the diagnostic information encoded by the RRL layer is largely, though not entirely, captured by the pathogen-derived classifier in the context of HIV status classification. This result is expected: the pathogen-derived classifier directly encodes the causal signal (viral antigen exposure), while the RRL signal is a downstream consequence of infection that is partially correlated with, but noisier than, the primary signal. The theoretical ceiling for integration gain is therefore constrained by the extent to which the RRL layer captures orthogonal classification-relevant variance — approximately 14% of cases were ranked differently between layers (Fig. 4A), providing the mechanistic basis for the observed, if modest, improvement.

These results do not diminish the value of the RRL layer; rather, they clarify its role. The RRL layer is better understood as an immune-state encoder than a diagnostic classifier. Its complementary information about host immunological architecture may become clinically relevant in contexts where pathogen-specific signals are diminished, such as early-stage infection prior to full seroconversion, post-treatment immune reconstitution, or monitoring of vaccine-induced versus infection-induced immune responses.

### 4.4 Repertoire Entropy and the Concentration Paradox in Chronic HIV Infection

The observation of significantly reduced Shannon entropy in HIV+ individuals (Mann–Whitney p = 0.012; Cliff’s δ = −0.30) across the RRL peptide binding distribution represents a conceptually important finding. HIV infection is clinically associated with hypergammaglobulinemia — markedly elevated total serum IgG — yet the present data indicate that this elevated antibody pool binds with greater concentration to a restricted subset of structural geometries. De Milito et al. demonstrated that hypergammaglobulinemia in HIV-1 infection arises in parallel to selective loss of antigen-specific antibodies and reduced memory B-cell frequencies (25), establishing a quantitative paradox: more total antibody, yet less specificity. The present entropy data resolve this paradox at the distributional level: increased total IgG is concentrated on a restricted set of structural binding geometries rather than distributed uniformly across the peptide landscape.

This pattern is mechanistically consistent with oligoclonal B-cell expansion following sustained antigen stimulation. The HIV Env-specific antibody response has been shown to be oligoclonal, consisting of expanded families derived from single B-cell clones as determined by V-D-J and V-J gene usage analysis (8). Chronic HIV viremia drives repeated cycles of B-cell activation, germinal center reactions, and clonal selection, ultimately producing a repertoire dominated by a limited number of expanded clones with specificities shaped by persistent antigenic pressure. Concurrently, naive B-cell diversity is depleted: naive B cells from HIV-1-infected patients show upregulated CD70 expression indicative of recent antigen stimulation and spontaneous IgG secretion (25), while memory B-cell subsets undergo progressive depletion that is not fully reversed by antiretroviral therapy (7, 30). Loss of this naive B-cell substrate removes the source of broad polyclonal responses, concentrating effective binding capacity among a smaller set of expanded specificities.

The differential performance of Gini index versus Shannon entropy in correlating with RRL classifier probability (r = 0.376 versus r = −0.278) provides additional interpretive resolution. The Gini coefficient is more sensitive to distributional inequality driven by a small number of dominant values, whereas Shannon entropy weights contributions across the full distribution. The stronger Gini correlation suggests that the classifier signal is driven primarily by the emergence of a few high-signal dominant peptides in HIV+ sera — consistent with clonal expansion producing high-affinity antibodies concentrated on a restricted set of structural targets. Moir et al. identified tissue-like memory B cells in HIV-viremic individuals with stunted Ig diversity and reduced replication histories, consistent with premature B-cell exhaustion (8), a phenomenon that would favor concentration rather than breadth of the antibody repertoire. This distinction favors a model of focal amplification over global suppression, with important implications for how immune repertoire remodeling is conceptualized in chronic infection.

The absence of significant correlations between classifier probability and either signal variance or top-peptide dominance (Fig. S6) is equally informative. These negative results indicate that the RRL classifier does not simply detect elevated total signal, signal spread, or domination by a single cross-reactive peptide. Rather, its diagnostic information derives from a distributional pattern involving multiple peptides simultaneously — a multi-peptide signature of immune topology. This guards against the interpretation that the RRL signal is a technical artifact of one cross-reactive peptide and instead supports a genuine multi-dimensional encoding of immune state.

### 4.5 A Conceptual Dual-Layer Model of Serology

Taken together, the results support a conceptual dual-layer model of serological information encoding in chronic viral infection. The first layer, represented by pathogen-derived peptides, encodes antigen specificity: it captures direct immunological memory of viral epitope exposure with high sensitivity and specificity, converging on the gp41 PID as the dominant diagnostic signal. This is consistent with the known immunodominance of this decoy region in natural HIV-1 infection across diverse patient populations (20). The second layer, represented by host-related combinatorial peptide signatures, encodes immune topology: it reflects the structural organization and distributional inequality of the global antibody repertoire as remodeled by chronic infection, and is a marker of systemic immunological state rather than antigen-specific memory.

These two layers are partially correlated because both are consequences of HIV infection, but they are not equivalent because they derive from different immunological processes operating at different timescales and with different degrees of individual variability. Antigen-specific memory is a near-universal consequence of seroconversion; immune repertoire remodeling is a continuous process shaped by viral load, treatment history, CD4+ T-cell dynamics, and host genetic factors. HIV-1 infection causes hypergammaglobulinemia, polyclonal activation, loss of memory B-cell subsets, B-cell exhaustion, and impaired humoral responses (7, 8), each of which contributes differently to the structural perturbations detectable by the RRL layer. This means the second layer may be more informative for tracking disease progression, treatment response, or vaccination effects than for initial diagnosis.

The methodological contribution of separating diagnostic ranking from immune-state encoding in the same dataset is a key aspect of this framework. Standard serological analyses evaluate classifiers exclusively by AUC, conflating these two dimensions. The present study demonstrates that distributional metrics — entropy, Gini index — applied to the raw peptide signal matrix can reveal immunological structure that is not captured by classifier probability alone. This multi-metric approach should be considered in the design and analysis of future peptide array-based serology studies.

### 4.6 Limitations and Future Directions

Several limitations of the present study warrant discussion. The cohort size (n = 105) is modest, and although strict nested cross-validation was implemented to prevent overfitting, independent external validation in a larger and more diverse cohort is necessary before clinical translation. The geographic diversity of the cohort (South Africa, Peru, Viet Nam) introduces potential confounders including HIV-1 subtype distribution, co-infection burden, and antiretroviral treatment prevalence, none of which were systematically controlled in the present analyses. Given that B-cell subset composition and hypergammaglobulinemia vary with viral load and treatment status (25, 30), a stratified sensitivity analysis by cohort and treatment status would be informative.

The absence of clinical metadata beyond HIV status represents a further limitation. CD4+ T-cell count and plasma viral load are the primary clinical correlates of HIV disease progression and immune deterioration. Testing whether Shannon entropy of the RRL binding distribution correlates with CD4+ count or viral load would directly test the hypothesis that the RRL layer encodes disease stage rather than merely infection status. Antiretroviral treatment (ART) status is a critical confounder: early ART has been shown to preserve resting memory B-cell subsets and normalize activated B-cell frequencies (31), and treated individuals might therefore show intermediate entropy values between untreated HIV+ and HIV− individuals. Resting memory B-cell depletion during chronic infection is not fully reverted upon successful ART (30), a finding which predicts persistent but attenuated entropy reduction in treated individuals — a testable hypothesis directly addressable with clinical metadata.

## Supporting information

readme

## Acknowledgments

The authors acknowledge financial support from the Ministerium für Wirtschaft, Arbeit und Tourismus Baden-Württemberg through the Invest BW innovation program.

## Conflict of Interest

C. von Bojnicic-Kninski and A. Nesterov-Mueller are cofounders of axxelera UG. All other authors declare no competing interests.

## Contributions

A.N.-M., N.S.-M., S.B. and A.D. conceived the project. A.N.-M and A.D. supervised the project. A.N.-M. developed the computational methodology and implemented the analysis pipeline. D.S. performed data preprocessing and statistical validation. C.v B.-K. and R.P. assisted with peptide chip design, sample incubation, and data interpretation. J.M. and H.B. contributed to the design of peptide libraries. J.G. performed all experimental work. All authors contributed to the interpretation of results. A.N.-M. and A.D. wrote the manuscript with input from all authors.

## Supplementary Material

**Figure S1:**
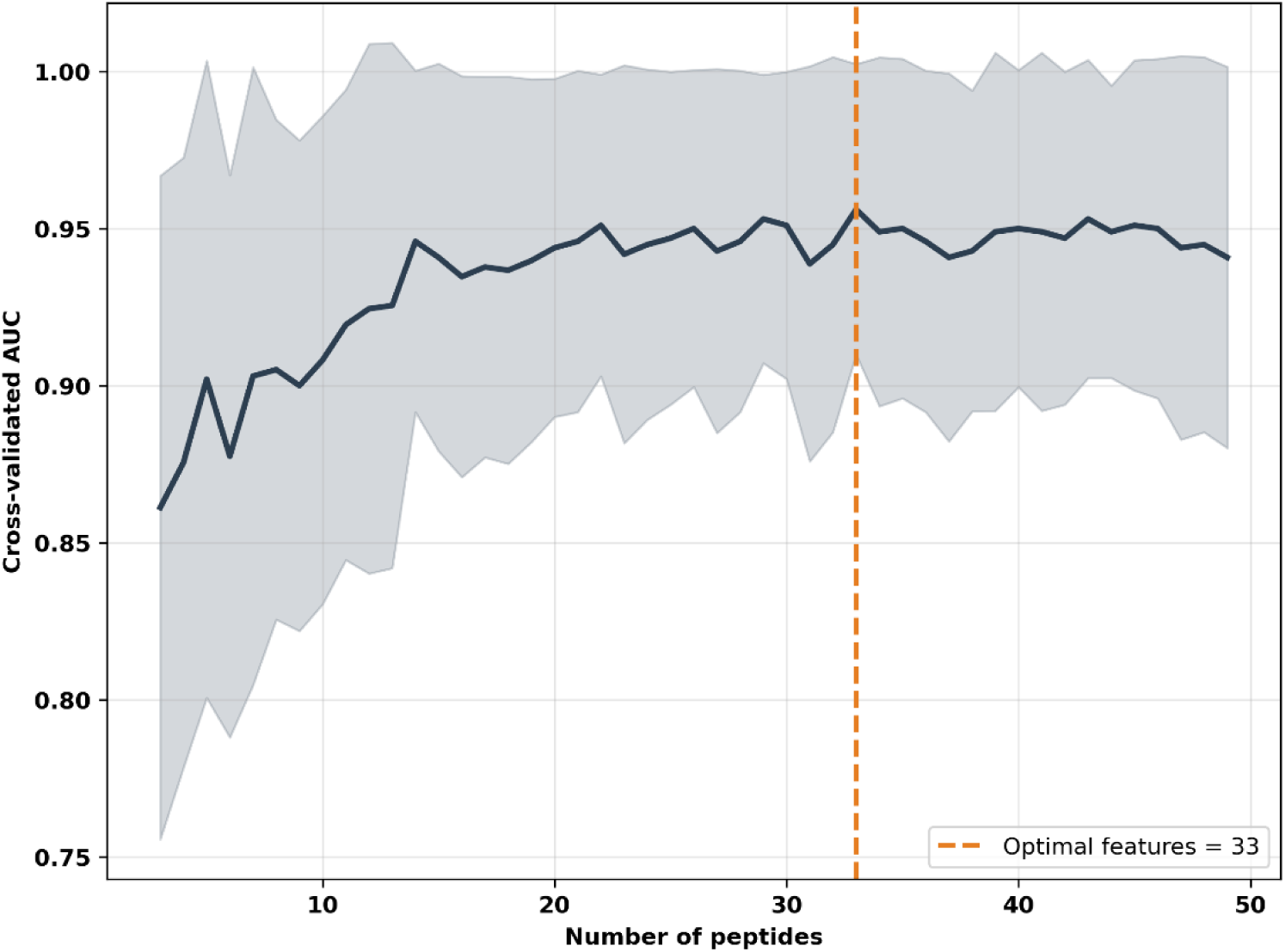
RFECV performance profile across peptide subset sizes. Recursive feature elimination with cross-validation (RFECV) was applied to the 49-peptide pathogen-derived panel using a Random Forest classifier (5-fold stratified cross-validation; scoring metric: AUC). The plot shows mean cross-validated AUC ± 1 s.d. as a function of the number of retained peptides. Model performance increases with feature count and reaches a broad maximum at ∼33 peptides (vertical dashed line), indicating convergence toward a compact feature subset with minimal additional performance gain beyond this range.

**Figure S2:**
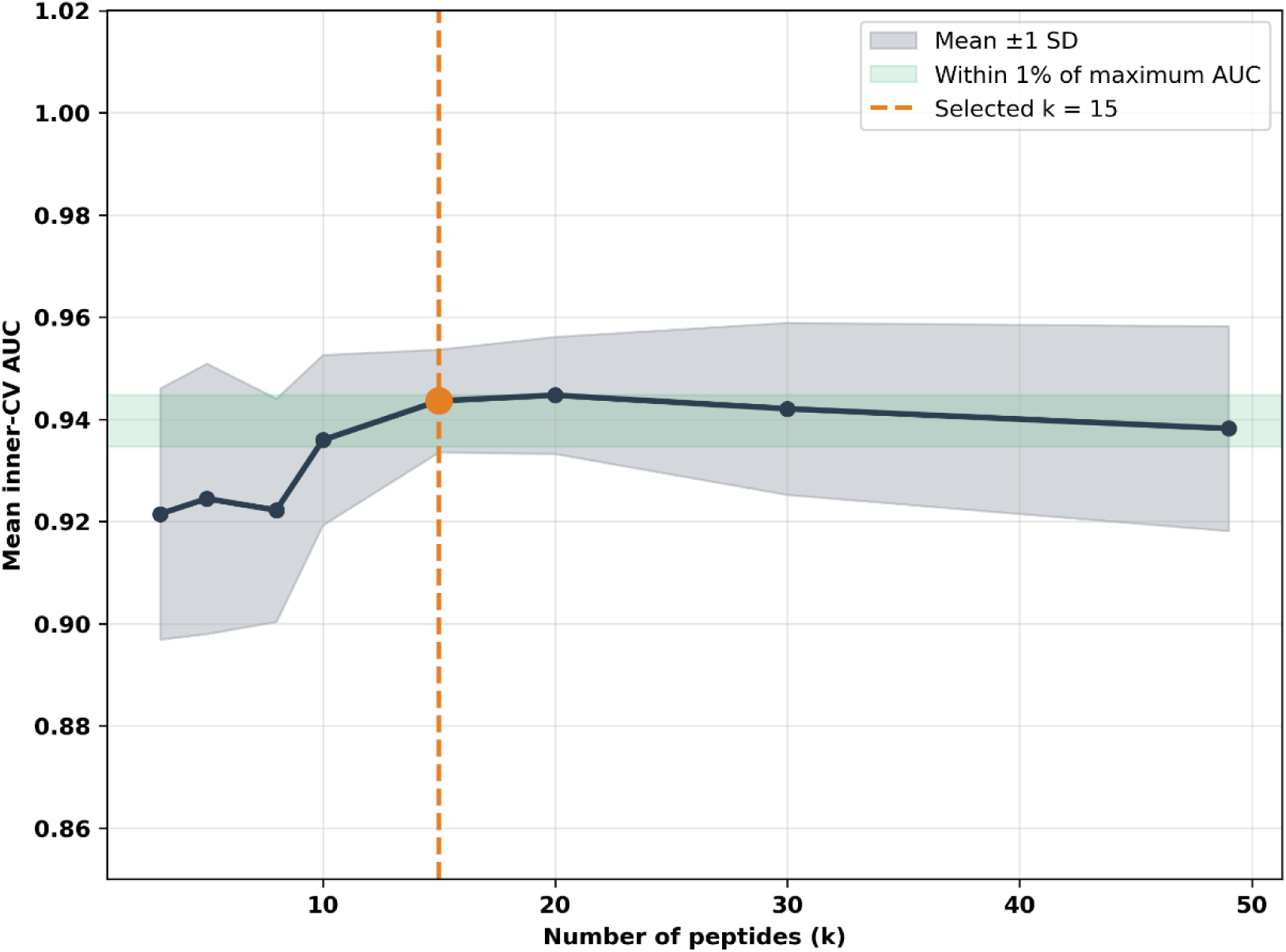
Performance plateau supports selection of a parsimonious 15-peptide panel. Inner cross-validated AUC (mean ± s.d., 5 folds) is plotted against the number of selected peptides (k) within the nested 5×5 cross-validation framework. Although the maximal AUC occurs at higher feature counts, performance remains within 1% of this maximum across a range of intermediate k values. The selected 15-peptide panel (vertical dashed line) lies within this performance equivalence interval, indicating that comparable diagnostic accuracy can be achieved with substantially fewer peptides, thereby supporting model parsimony without measurable loss of predictive performance.

**Figure S3:**
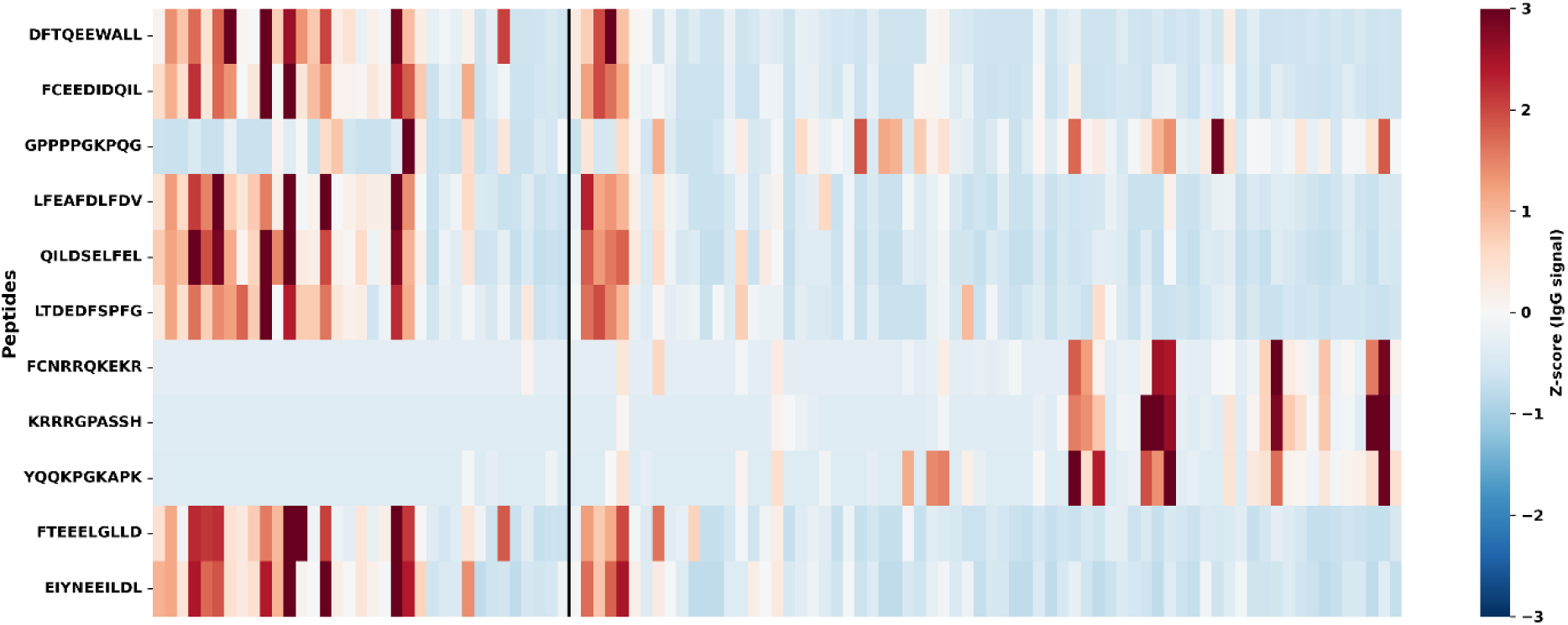
Z-scored IgG reactivity across stable peptides from the host-related layer. Heatmap of Z-scored IgG signal intensities for peptides selected in ≥3/5 outer cross-validation folds. Signals were standardized per peptide across all samples (n = 105) and clipped at ±3 standard deviations. Samples are grouped by HIV status and ordered within each group by predicted probability from the RRL classifier. The vertical black line marks separation between HIV+ and HIV− individuals. Color scale represents standardized IgG signal intensity.

**Figure S4:**
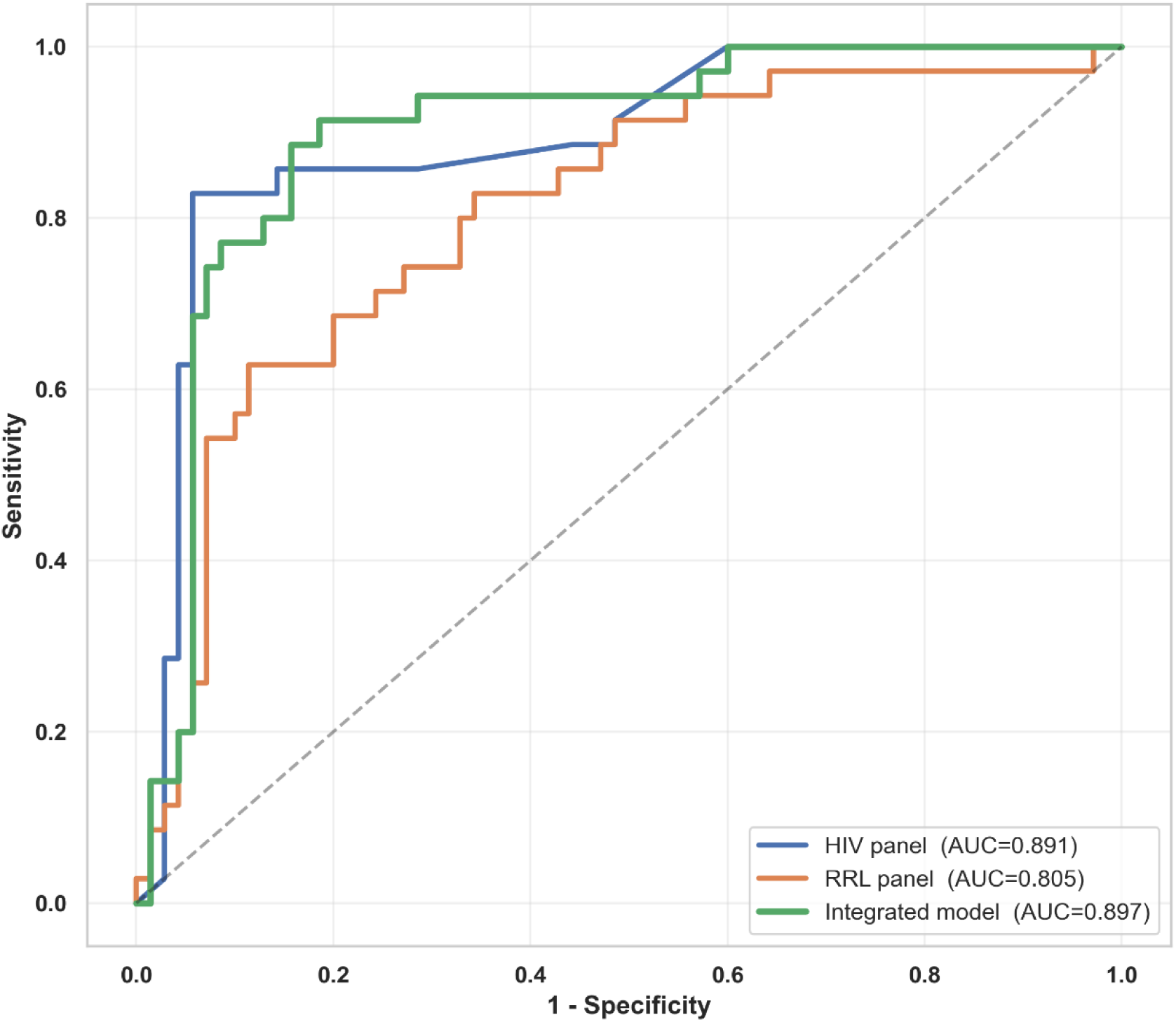
ROC comparison of individual and integrated serological classifiers. RO curves for the pathogen-derived HIV peptide panel, the RRL panel, and a logistic regression meta-model integrating out-of-fold probabilities from both classifiers (nested cross-validation; n = 105). While the integrated model achieved a marginal increase in AUC (0.897) compared with the HIV panel alone (0.891), the near-overlapping ROC profiles indicate that most diagnostic discrimination is driven by the pathogen-derived signal.

**Figure S5:**
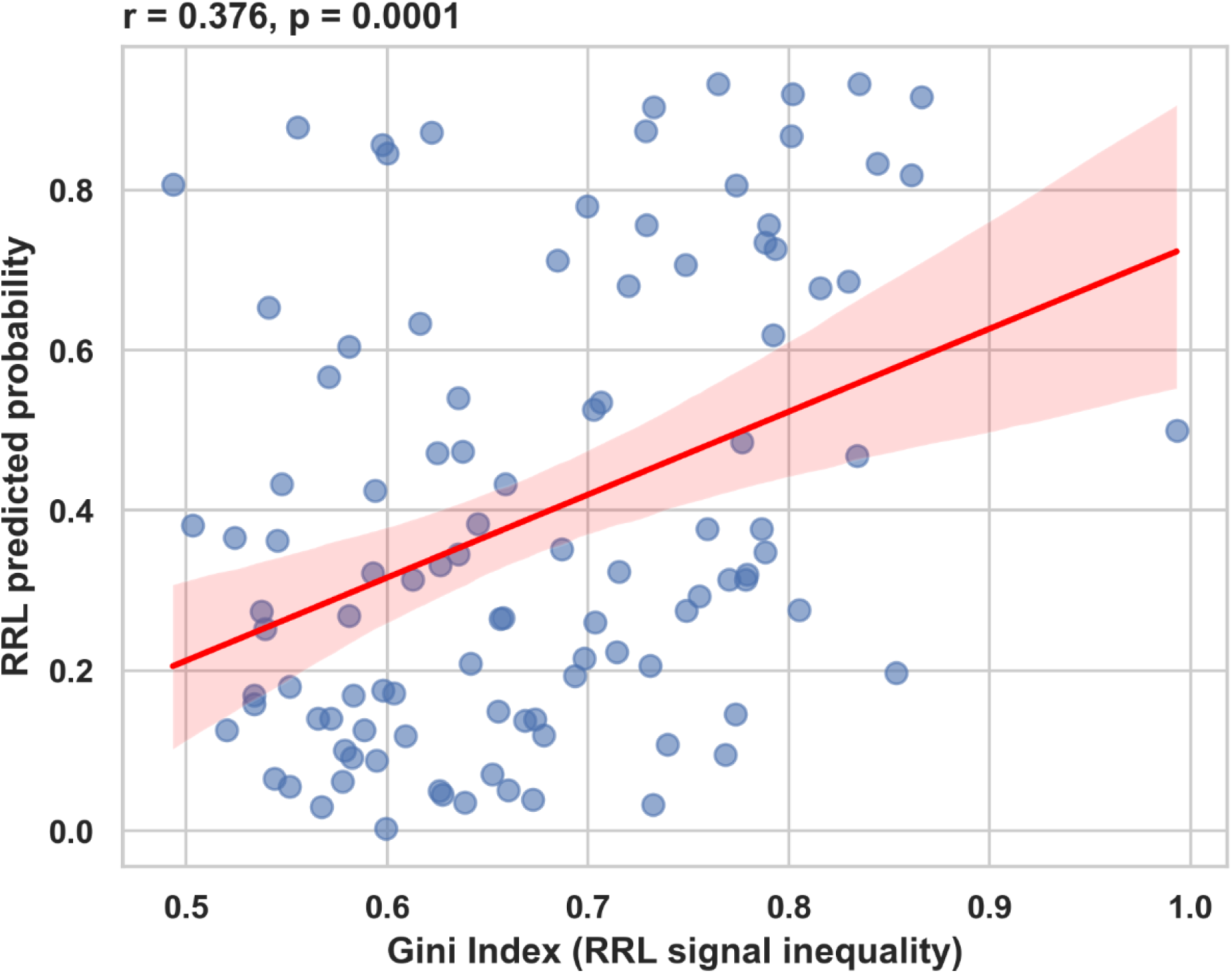
Gini index correlates with RRL classifier output. Relationship between the Gini index (per-sample signal inequality across the RRL peptide set) and RRL-predicted probability. Pearson correlation coefficient and corresponding p-value are indicated in the panel. The regression line denotes the best linear fit.

**Figure S6:**
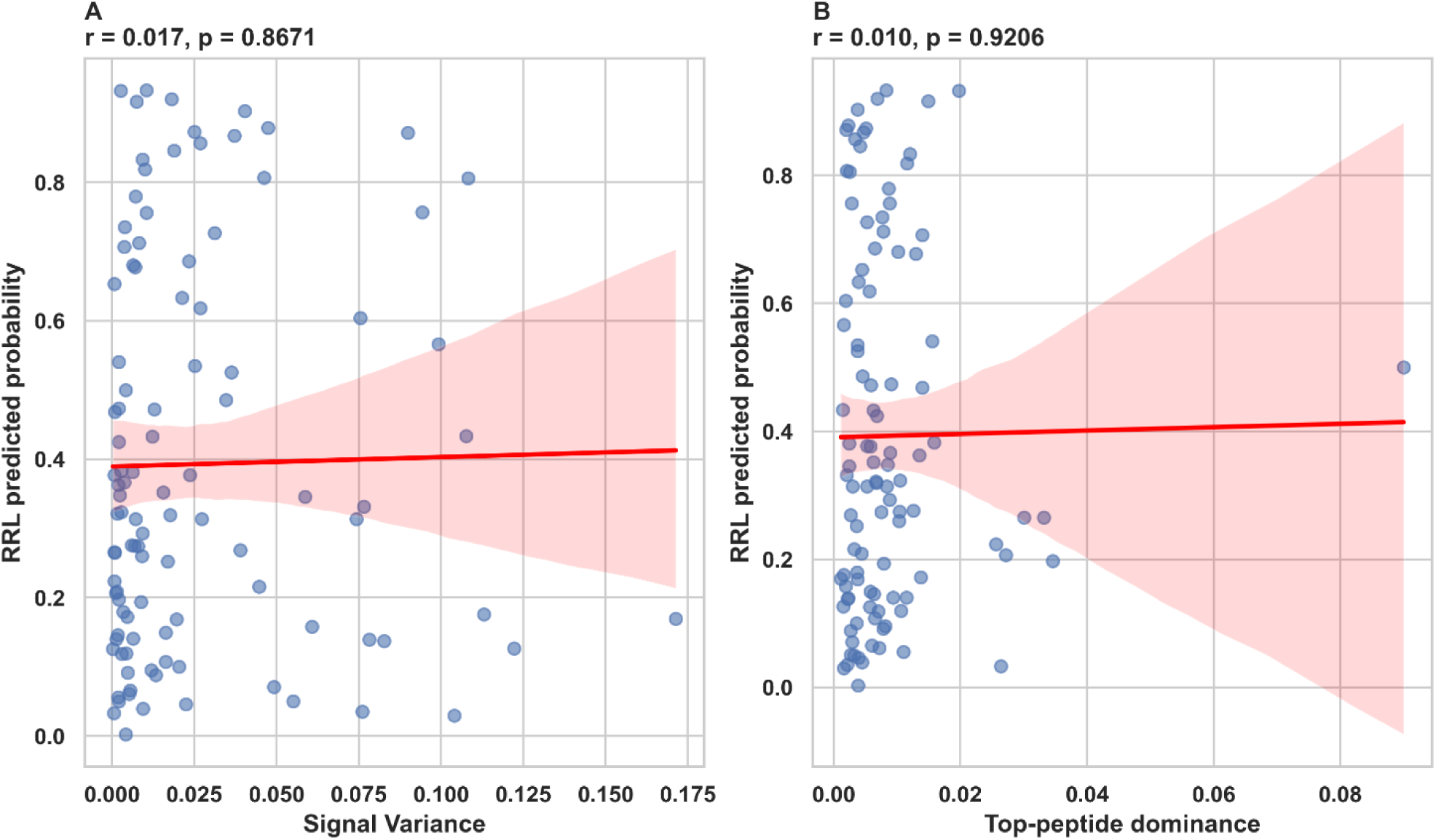
Variance and top-peptide dominance do not correlate with RRL classifier output. (A) Scatter plot of per-sample signal variance versus RRL-predicted probability (n = 105). Pearson correlation coefficient and p-value are indicated in the panel. (B) Scatter plot of top-peptide dominance versus RRL-predicted probability. No significant association was detected.

### Supplementary Methods S1. Selection of HIV-1 peptides

#### S1.1 Source databases and protein targets

Peptides representing HIV-1 were derived from antibody-epitope alignments of the p24 major core-capsid protein (*p24_ab_align.fasta*, downloaded on 28 June 2023) and of the envelope protein Env (*Env_ab_align.fasta*, downloaded on 29 June 2023), both obtained from the Los Alamos National Laboratory (LANL) HIV Immunology Database (32, 33). Inspection of the LANL antibody-epitope density map (34) identified p24 and the more conserved gp41 subunit of Env as the two regions carrying the highest density of mapped linear antibody epitopes. Consequently, peptide library design was restricted to p24 and gp41.

The LANL p24 alignment corresponds to residues 133–263 of the Gag polyprotein of the HXB2 reference strain (GenBank nucleotide accession *K03455*; protein accession *AAB50258.1*). The Gag polyprotein is proteolytically processed at five sites into the matrix protein p17, the p24 capsid protein, the p7 nucleocapsid, p6 and two short SP1/SP2 peptides (35); p24 subsequently assembles into the hexameric and pentameric rings that form the mature capsid shell (36). The Env alignment corresponds to the 856-residue envelope polyprotein precursor gp160 (protein accession *AAB50262.1*), in which residues 1–30 form the signal peptide, 31–511 the surface glycoprotein gp120 and 512–856 the transmembrane subunit gp41. After signal-peptide removal and furin cleavage between residues 511/512, the trimeric gp120–gp41 envelope complex is assembled (37, 38).

#### S1.2 Selection of regions of interest

Regions harboring linear antibody epitopes in p24 and Env were deduced from the LANL antibody-epitope density map and from the LANL “Linear antibody epitopes” summary table (*lalnl_HIV_ab_summary.csv*, downloaded on 28 June 2023) (34). For p24, an immunodominant linear epitope originally mapped to residues 111–125 by Graham *et al.* (39) was not selected, as this region is among the least conserved in the protein and therefore unsuitable for pan-clade diagnostic purposes. Janvier *et al.* (40) subsequently reported that p24 regions 46–60 and 156–170 reacted with 40–45 % of HIV-positive sera and probably form a discontinuous, two-loop conformational epitope; both regions were therefore retained in the present design. Diagnostic utility of the p24 N- and C-terminal 30-mer regions and of chimeric constructs thereof reported by Hernández *et al.* (41) was further reason to include the N- and C-terminal regions.

Stephenson *et al.* (42) designed a 15-mer tiling peptide microarray with 14-residue overlap covering large parts of the HIV-1 LANL proteome (6,564 peptides; p24, 264 peptides, 86.2 % sequence coverage; gp120, 2,672 peptides, 50.2 %; gp41, 1,210 peptides, 65.5 %). In that study, the main responding gp41 regions mapped to Env positions 560–620 and 640–690, in agreement with the LANL linear-epitope density map, and both regions were therefore included here. Li *et al.* (43) tested long peptides spanning three major linear-epitope regions in p24 (45–93, 148–187, 188–227) and in gp41 (560–616, 639–686). Chinese sera did not react with the p24 peptides but reacted strongly with the gp41 560–616 peptide. In view of these findings, the gp41 region 560–616 was considered the single most informative diagnostic target. Consequently, the final regions retained for library generation were p24 residues 1–30, 44–60, 64–135, 140–190 and 202–231, and gp41 (Env numbering) residues 560–616 and 639–686.

#### S1.3 In-silico peptide generation

To capture HIV-1 diversity across these regions while keeping the library below 10,000 peptides, the LANL 2021 “Subtype Reference Alignments” of the HIV Sequence Compendium (44) were used. These alignments contain 198 HIV-1 sequences, covering the HXB2 reference strain, four representatives of each clade and major circulating recombinant forms, and are annotated for all HIV-1 proteins in the accompanying LANL protein file *hiv1prot.pdf* (44). A Perl script was used to extract 12-mer peptides, overlapping by 11 residues, from the p24 and Env alignment files, stepping through each aligned sequence at the coordinates of the selected regions. Peptides containing the placeholder “X” (undefined residue) or identical to a peptide previously extracted from an earlier sequence were discarded. This procedure yielded 4,537 unique p24-derived peptides and 5,160 unique gp41-derived peptides, for a combined set of 9,697 12-mer sequences.

#### S1.4 Selection of strongly reacting peptides from test arrays

From the 9,697 candidate peptides, a subset was chosen on the basis of empirical reactivity across two independent test-array experiments. Peptides were retained if they produced a strong signal with HIV-positive tuberculosis-panel sera and a low background with HIV-negative sera in both runs. Forty-six peptides met these criteria: 13 p24-derived peptides and 33 gp41-derived peptides (Supplementary Table S1). Within the p24 set, region 64–135 contributed seven peptides, region 140–190 contributed five peptides, and the C-terminal region 202–231 contributed one peptide. Six of the 13 p24 peptides cover the cyclophilin-A (CypA)–binding region (HXB2 annotation 72–108); three of these span the full CypA-binding loop at residues 85–93 (HIV_p24_2116, HIV_p24_2720 and HIV_p24_2750, all covering positions 82–93). CypA binding is essential for nuclear import and reverse transcription, and polymorphisms in this loop modulate viral resistance against the interferon-induced host defense protein MxB (45).

Within the 33 gp41-derived peptides, 25 mapped to the major immunodominant region in Env (LANL annotation 588–607). This segment lies within the highly flexible 46-residue linker that connects the two conserved helical-bundle regions HR1 and HR2. In the post-fusion Env structure, the three N36 helices of HR1 (residues 546–581) form the inner stalk of a six-helix bundle, which is surrounded by the three C34 helices of HR2 (residues 628–661) (38). The six selected peptides in region 639–667 overlap the C-terminal half of the C34 helix. The most C-terminal selected peptide (HIV_env_4258, HXB2 residues 656–667) covers the first eight residues of the membrane-proximal external region (MPER, 661–683), which is a primary target of rare but exceptionally broadly neutralising antibodies (46).

#### S1.5 Randomized control peptides

To discriminate sequence-specific epitope recognition from binding that depends solely on amino-acid composition, Randomized control peptides were generated for three representative high-intensity peptides. Randomized controls contain the same amino-acid composition as their template peptide but with residue positions randomly permuted by a Perl script (listed at the bottom of Supplementary Table S1). Concordant strong binding of a serum to both a template and its scrambled counterpart indicates non-specific, composition-driven interaction, whereas binding restricted to the template indicates sequence-specific epitope recognition.

**Supplementary Table S1.**
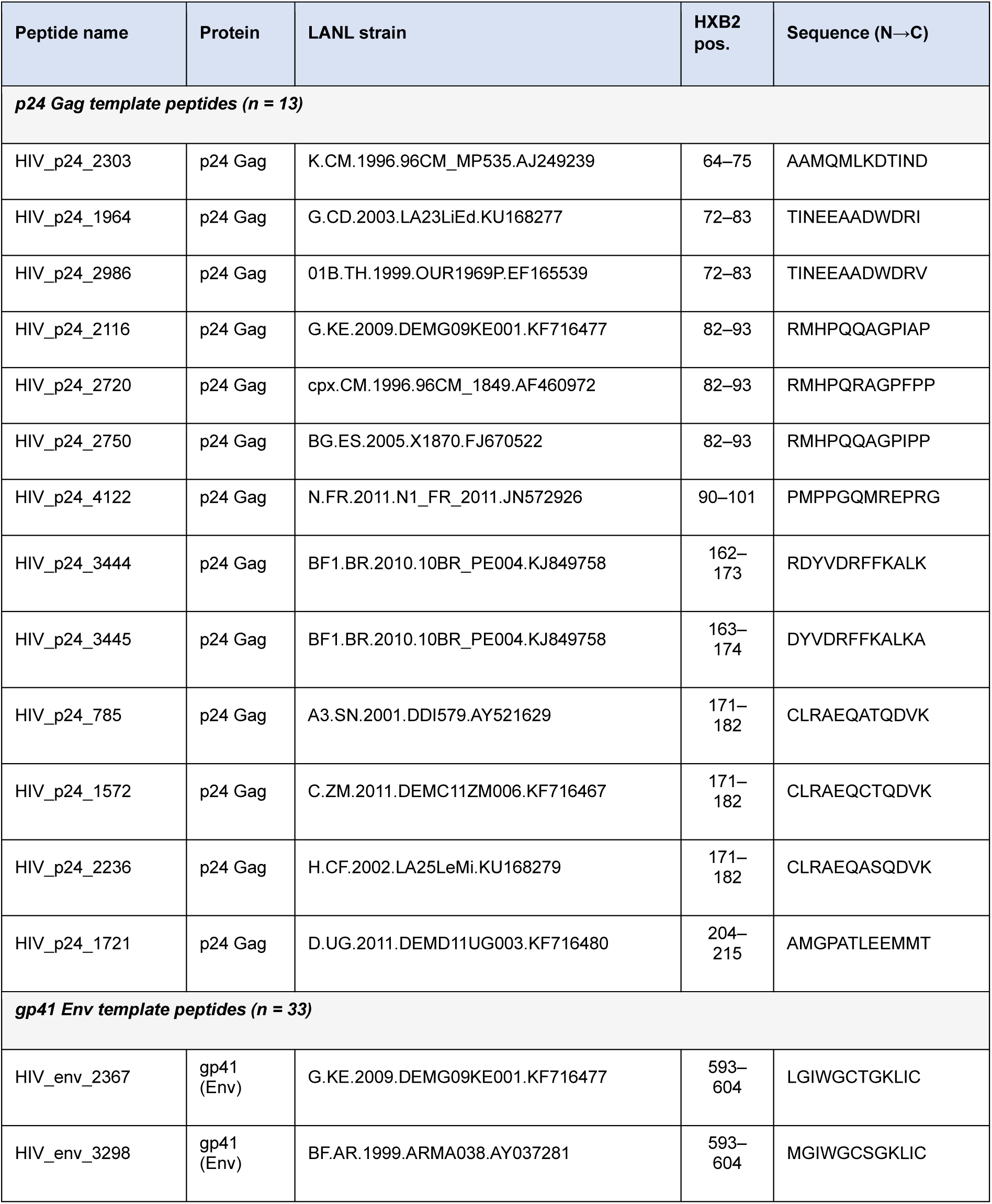
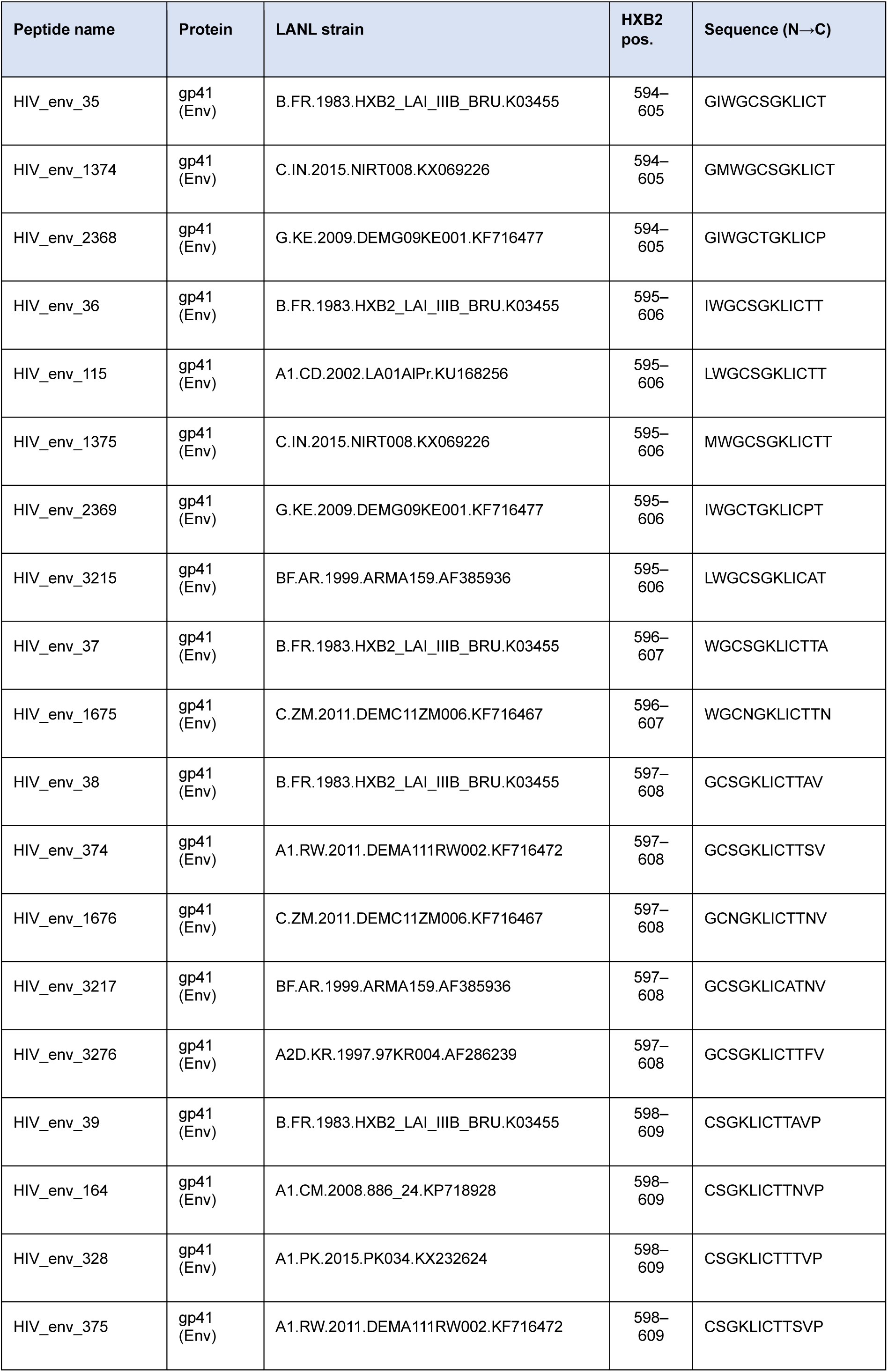
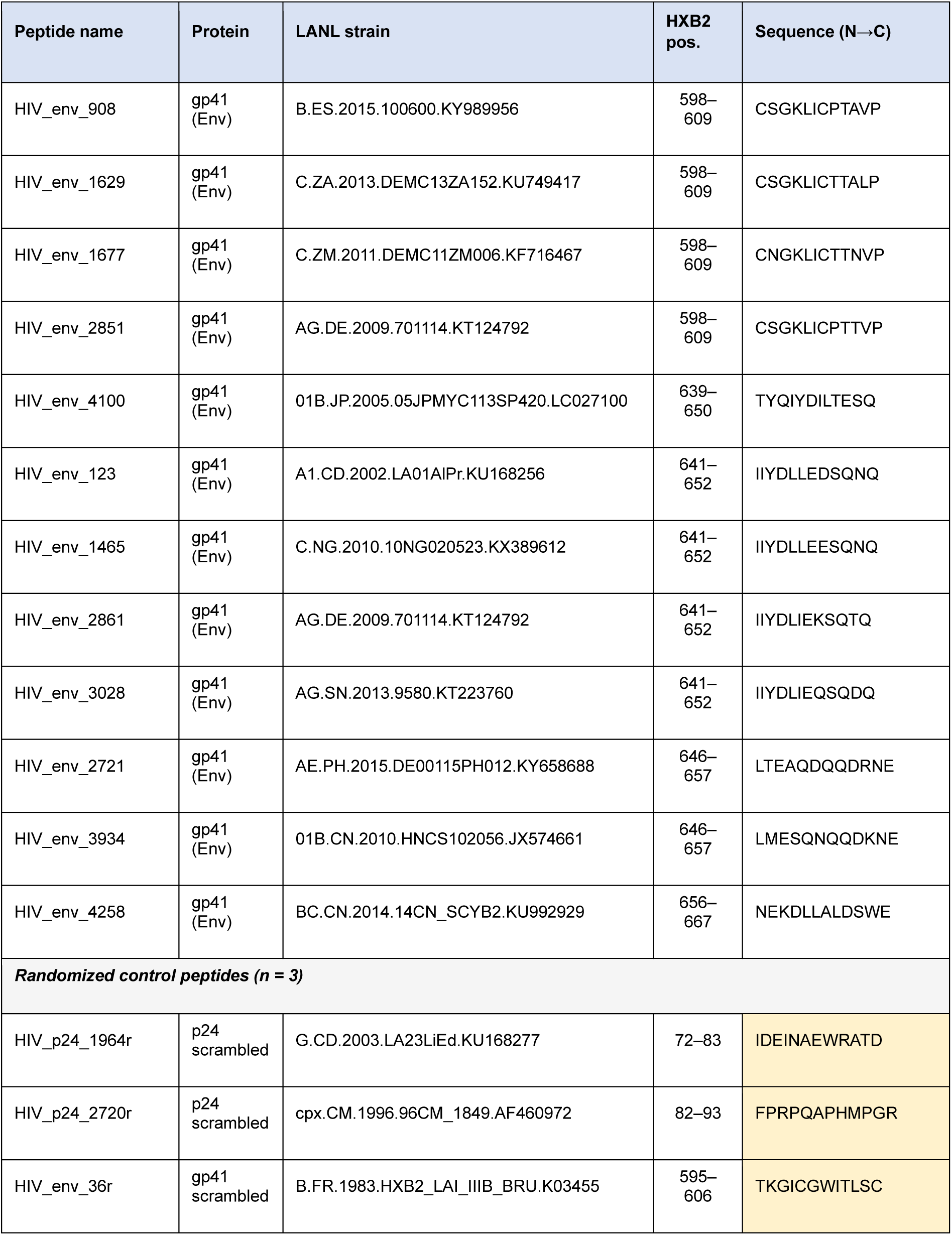
HIV-1 peptide panel used for nested cross-validation modelling (*n* = 49 peptides).

