## Supplementary material for "Dual-Layer Serological Encoding: Integrating Pathogen-Derived and Host-Reactive Peptide Signatures Reveals Complementary Dimensions of HIV Immune Response": readme

# Supplementary Code and Data

## Dual-Layer Serological Encoding: Integrating Pathogen-Derived and Host-Reactive Peptide Signatures Reveals Complementary Dimensions of HIV Immune Response

This repository contains the raw peptide microarray data and the Python/Jupyter analysis pipeline
required to reproduce every numerical result and figure in the manuscript. The analysis evaluates a
dual-layer serological framework combining a pathogen-derived HIV peptide panel (49 peptides) with a
host-related resemblance-ranking peptide library (RRL; 2,087 peptides) in a cohort of 105 serum
samples (35 HIV+, 70 HIV−) from South Africa, Peru and Viet Nam.

---

## 1. File inventory

| File | Type | Purpose |
|---|---|---|
| `results_HIV_final.pkl` | Data (pickle) | Raw IgG fluorescence signals, 49 HIV-derived peptides × 105 samples |
| `results_RRL_final.pkl` | Data (pickle) | Raw IgG fluorescence signals, 2,087 host-related RRL peptides × 105 samples |
| `HIV_RF.ipynb` | Analysis notebook | Layer 1 — Random Forest classification on the HIV panel (nested 5×5 CV) |
| `RRL_RF.ipynb` | Analysis notebook | Layer 2 — Random Forest classification on the RRL library (nested 5×5 CV) |
| `Covariance.ipynb` | Analysis notebook | Meta-model integration, orthogonality analysis, entropy / Gini metrics |

---

## 2. Data description

### 2.1 `results_HIV_final.pkl`

Python pickle file containing a `dict` with 105 entries, one per serum sample. Keys are sample
identifiers of the form `S1`, `S2`, …, `S116` (gaps reflect samples excluded during QC).

Each value is a `pandas.DataFrame` with the following schema:

| Column | dtype | Description |
|---|---|---|
| `ID` | str | Peptide amino-acid sequence (49 rows per sample) |
| `MedianValue` | float | Median IgG fluorescence intensity across replicate spots |
| `StdDevValue` | float | Standard deviation of fluorescence across replicate spots |
| `ClasPep` | float (0/1) | Internal peptide-level reactivity flag |
| `SampleID` | str | Sample identifier (matches the dictionary key) |

The 49 peptides comprise 34 gp41 envelope peptides and 15 p24 Gag peptides.

### 2.2 `results_RRL_final.pkl`

Python pickle file containing a `dict` with 105 entries (same sample keys as above).

Each value is a `pandas.DataFrame` with:

| Column | dtype | Description |
|---|---|---|
| `ID` | str | Peptide amino-acid sequence (2,087 rows per sample) |
| `MedianValue` | float | Median IgG fluorescence intensity across replicate spots |
| `StdDevValue` | float | Standard deviation of fluorescence across replicate spots |

The RRL peptide set was generated according to the resemblance-ranking design described in
Jenne et al., *Int. J. Mol. Sci.* **23**, 3515 (2022).

### 2.3 Sample labels

HIV status (the binary classification endpoint) is not embedded in the pickle files. Labels
(`HIV+ = 1`, `HIV− = 0`) are supplied by the accompanying metadata table referenced inside the
notebooks as `df_samples_all.xls`. This file is available from the corresponding author on
reasonable request, subject to the data-sharing conditions of the original sample biobanks.

---

## 3. Reproducing the analysis

The notebooks are designed to be run in the following order. Each notebook exports intermediate
files that are consumed by the next step.

### Step 1 — `HIV_RF.ipynb` (Layer 1)

**Input:** `results_HIV_final.pkl`, sample metadata
**Pipeline:**
- Assembly of the feature matrix (samples × 49 peptides)
- Nested stratified cross-validation (outer 5-fold, inner 5-fold)
- Feature ranking by Random Forest Gini importance (inside each outer training fold)
- Adaptive *k*-selection (smallest *k* within 1 % of best inner-fold AUC)
- Random Forest classifier: `n_estimators = 500`, `class_weight = "balanced"`
- Recursive feature elimination with cross-validation (RFECV)
- Permutation test (100 permutations)

**Outputs:** pooled out-of-fold (OOF) probabilities, stability frequencies, ROC curve, z-scored
reactivity heatmap.
**Reproduces:** Figure 1, Figure 2, Supplementary Figure S1, Supplementary Figure S2.

### Step 2 — `RRL_RF.ipynb` (Layer 2)

**Input:** `results_RRL_final.pkl`, sample metadata
**Pipeline:** identical nested 5×5 CV framework as Step 1, adapted to the high-dimensional
setting (*p* ≈ 2,087, *n* = 105).

**Outputs:** OOF probabilities, stability frequencies, ROC curve, z-scored reactivity heatmap.
**Reproduces:** Figure 3, Supplementary Figure S3.

### Step 3 — `Covariance.ipynb` (integration and immune-topology analysis)

**Input:** the OOF probability files exported by Steps 1 and 2
(`HIV_RF_n105_oof_predictions.csv`, `RRL_RF_n105_oof_predictions.csv`), plus
`results_RRL_final.pkl` for distributional metrics.

**Pipeline:**
- Pearson correlation of OOF probabilities with 10,000-iteration bootstrap 95 % confidence interval
- Logistic-regression meta-model using `[P_RRL, P_HIV, P_RRL × P_HIV]` as features, trained under
  stratified 5-fold outer CV with inner 3-fold `GridSearchCV` over `C ∈ {0.01, 0.1, 1, 5, 10}`
- Per-sample Shannon entropy, Gini index, signal variance and top-peptide dominance over the
  2,087 RRL peptides
- Mann–Whitney *U* test between HIV+ and HIV− groups; Cliff's δ as effect size
- Pearson and Spearman correlations between distributional metrics and RRL classifier probability

**Reproduces:** Figure 4, Figure 5, Supplementary Figures S4, S5, S6.

---

## 4. Software environment

All analyses were performed in Python 3.10+. The notebooks depend on the following packages
(versions tested during manuscript preparation; newer versions are expected to be compatible):

- `numpy` ≥ 1.23
- `pandas` ≥ 1.5
- `scikit-learn` ≥ 1.2
- `scipy` ≥ 1.10
- `matplotlib` ≥ 3.6
- `seaborn` ≥ 0.12

Install with:

```bash
pip install numpy pandas scikit-learn scipy matplotlib seaborn jupyter
```

All random operations use a fixed seed (`random_state = 42` / `RS = 42`) so that cross-validation
folds, Random Forest bootstrap samples and bootstrap confidence intervals are exactly reproducible.

Loading the pickle files:

```python
import pickle
with open("results_HIV_final.pkl", "rb") as f:
    hiv_data = pickle.load(f)        # dict[sample_id -> DataFrame (49, 5)]
with open("results_RRL_final.pkl", "rb") as f:
    rrl_data = pickle.load(f)        # dict[sample_id -> DataFrame (2087, 3)]
```

---

## 5. Notes for reviewers

- All preprocessing, standardization, feature ranking and hyperparameter tuning are strictly
  nested inside the outer cross-validation loop. No sample-level information from a held-out
  outer fold is used at any stage of model training on the corresponding training fold.
- Out-of-fold probabilities are pooled across the five outer folds before computing AUC and
  before entering the meta-model, which prevents optimistic bias from in-fold predictions.
- The meta-model itself is evaluated under a separate nested 5×5 scheme, so the reported
  integrated AUC of 0.897 is also an out-of-fold estimate.

---

## 6. Contact

Questions regarding the code or the raw data should be directed to the corresponding author of
the manuscript.
