## Supplementary material for "Dual-Layer Serological Encoding: Integrating Pathogen-Derived and Host-Reactive Peptide Signatures Reveals Complementary Dimensions of HIV Immune Response": Code and Data: Covariance.ipynb

```
# %% [code]
"""
=========================================================
 ORTHOGONALITY OF SEROLOGICAL LAYERS
 RRL vs HIV peptide panels (OOF probabilities)
=========================================================
"""

import numpy as np
import pandas as pd
import matplotlib.pyplot as plt
import seaborn as sns
from scipy.stats import pearsonr
from sklearn.utils import resample

# ─────────────────────────────────────────────
# PATHS
# ─────────────────────────────────────────────
DATA_PATH = r"D:\Users\ha6057\Desktop\HIV_AI\Files for Analysis with Claud"

RRL_FILE = DATA_PATH + r"\RRL_RF_n105_oof_predictions.csv"
HIV_FILE = DATA_PATH + r"\HIV_RF_n105_oof_predictions.csv"

# ─────────────────────────────────────────────
# LOAD
# ─────────────────────────────────────────────
rrl = pd.read_csv(RRL_FILE)
hiv = pd.read_csv(HIV_FILE)

df = rrl.merge(hiv, on="sample", suffixes=("_RRL", "_HIV"))

print("\nMerged samples:", len(df))

# ─────────────────────────────────────────────
# PEARSON CORRELATION
# ─────────────────────────────────────────────
r, p = pearsonr(df["oof_prob_RRL"], df["oof_prob_HIV"])

print(f"\nPearson r = {r:.4f}")
print(f"p-value   = {p:.6f}")

# ─────────────────────────────────────────────
# BOOTSTRAP CI
# ─────────────────────────────────────────────
n_boot = 10000
boot_r = []

for _ in range(n_boot):
    sample_df = resample(df)
    rb, _ = pearsonr(sample_df["oof_prob_RRL"],
                     sample_df["oof_prob_HIV"])
    boot_r.append(rb)

ci_low  = np.percentile(boot_r, 2.5)
ci_high = np.percentile(boot_r, 97.5)

print(f"95% Bootstrap CI: [{ci_low:.4f}, {ci_high:.4f}]")


# %% [code]
"""
=========================================================
 IMPROVED META-MODEL (SAFE, REGULARIZED)
 RRL + HIV probabilities
=========================================================
"""

from sklearn.linear_model import LogisticRegression
from sklearn.model_selection import StratifiedKFold, GridSearchCV
from sklearn.preprocessing import StandardScaler
from sklearn.pipeline import Pipeline
from sklearn.metrics import roc_auc_score
import numpy as np
import pandas as pd

# ─────────────────────────────────────────────
# LOAD
# ─────────────────────────────────────────────
DATA_PATH = r"D:\Users\ha6057\Desktop\HIV_AI\Files for Analysis with Claud"

rrl = pd.read_csv(DATA_PATH + r"\RRL_RF_n105_oof_predictions.csv")
hiv = pd.read_csv(DATA_PATH + r"\HIV_RF_n105_oof_predictions.csv")

df = rrl.merge(hiv, on="sample", suffixes=("_RRL", "_HIV"))

X = np.column_stack([
    df["oof_prob_RRL"],
    df["oof_prob_HIV"]
])

# interaction term
interaction = (X[:,0] * X[:,1]).reshape(-1,1)
X = np.hstack([X, interaction])

y = df["true_label_RRL"].values

# ─────────────────────────────────────────────
# META MODEL
# ─────────────────────────────────────────────
outer_cv = StratifiedKFold(n_splits=5, shuffle=True, random_state=42)
meta_probs = np.zeros(len(y))

for train_idx, test_idx in outer_cv.split(X, y):

    X_tr, X_te = X[train_idx], X[test_idx]
    y_tr = y[train_idx]

    pipe = Pipeline([
        ("scaler", StandardScaler()),
        ("clf", LogisticRegression(
            solver="lbfgs",
            max_iter=2000
        ))
    ])

    param_grid = {
        "clf__C": [0.01, 0.1, 1, 5, 10]
    }

    grid = GridSearchCV(
        pipe,
        param_grid,
        cv=3,
        scoring="roc_auc"
    )

    grid.fit(X_tr, y_tr)

    meta_probs[test_idx] = grid.best_estimator_.predict_proba(X_te)[:,1]

# ─────────────────────────────────────────────
# PERFORMANCE
# ─────────────────────────────────────────────
auc_meta = roc_auc_score(y, meta_probs)
auc_rrl  = roc_auc_score(y, df["oof_prob_RRL"])
auc_hiv  = roc_auc_score(y, df["oof_prob_HIV"])

print("\nAUC RRL :", round(auc_rrl,4))
print("AUC HIV :", round(auc_hiv,4))
print("AUC META:", round(auc_meta,4))
print("Improvement:", round(auc_meta - max(auc_rrl, auc_hiv),4))

# %% [code]
# ═══════════════════════════════════════════════════════
# SUPPLEMENTARY FIGURE — Integration ROC Overlay
# ═══════════════════════════════════════════════════════

import matplotlib.pyplot as plt
from sklearn.metrics import roc_curve
from sklearn.utils import resample
import numpy as np
import os

OUT_FILE = os.path.join(DATA_PATH, "Supplementary_Fig4_Integration_ROC.png")

# ------------------------------------------------------
# ROC CURVES
# ------------------------------------------------------
fpr_rrl, tpr_rrl, _   = roc_curve(y, df["oof_prob_RRL"])
fpr_hiv, tpr_hiv, _   = roc_curve(y, df["oof_prob_HIV"])
fpr_meta, tpr_meta, _ = roc_curve(y, meta_probs)

# ------------------------------------------------------
# BOOTSTRAP ΔAUC (META − HIV)
# ------------------------------------------------------
n_boot = 10000
delta_aucs = []

for _ in range(n_boot):
    sample_idx = resample(np.arange(len(y)))
    yb = y[sample_idx]
    
    auc_h = roc_auc_score(yb, df["oof_prob_HIV"].values[sample_idx])
    auc_m = roc_auc_score(yb, meta_probs[sample_idx])
    
    delta_aucs.append(auc_m - auc_h)

delta_low  = np.percentile(delta_aucs, 2.5)
delta_high = np.percentile(delta_aucs, 97.5)

# ------------------------------------------------------
# PLOT
# ------------------------------------------------------
fig, ax = plt.subplots(figsize=(8,7), dpi=300)

plt.plot(fpr_hiv, tpr_hiv,
         label=f"HIV panel  (AUC={auc_hiv:.3f})",
         linewidth=2.5)

plt.plot(fpr_rrl, tpr_rrl,
         label=f"RRL panel  (AUC={auc_rrl:.3f})",
         linewidth=2.5)

plt.plot(fpr_meta, tpr_meta,
         label=f"Integrated model  (AUC={auc_meta:.3f})",
         linewidth=3)

plt.plot([0,1],[0,1],'k--',alpha=0.4)

plt.xlabel("1 - Specificity", fontweight="bold")
plt.ylabel("Sensitivity", fontweight="bold")

#plt.title(
#    f"Probability-Level Integration\n"
#    f"ΔAUC = {auc_meta - auc_hiv:.3f}  "
#    f"(95% CI [{delta_low:.3f}, {delta_high:.3f}])",
#    fontweight="bold"
#)

        
plt.legend()
plt.grid(alpha=0.3)
plt.tight_layout()
# Make tick labels bold
# Make tick labels bold
for tick in ax.get_xticklabels():
    tick.set_fontweight("bold")
for tick in ax.get_yticklabels():
    tick.set_fontweight("bold")
    
plt.savefig(OUT_FILE, bbox_inches="tight")
plt.show()

print("Supplementary integration ROC figure 4 saved.")

# %% [code]
# ═══════════════════════════════════════════════
# FIGURE 4 — Partial Orthogonality (A + B)
# ═══════════════════════════════════════════════

import matplotlib.pyplot as plt
import seaborn as sns
import numpy as np
from scipy.stats import pearsonr
from sklearn.utils import resample
import os

OUT_FILE = os.path.join(DATA_PATH, "Figure4_Orthogonality.png")

# ------------------------------------------------------
# Pearson correlation
# ------------------------------------------------------
r, p = pearsonr(df["oof_prob_RRL"], df["oof_prob_HIV"])

# Bootstrap CI
n_boot = 10000
boot_r = []

for _ in range(n_boot):
    sample_df = resample(df)
    rb, _ = pearsonr(sample_df["oof_prob_RRL"],
                     sample_df["oof_prob_HIV"])
    boot_r.append(rb)

ci_low  = np.percentile(boot_r, 2.5)
ci_high = np.percentile(boot_r, 97.5)

r2 = r**2

# ------------------------------------------------------
# FIGURE
# ------------------------------------------------------
sns.set_style("whitegrid")

fig, axes = plt.subplots(2, 1, figsize=(8, 10), dpi=300)

# ═══════════════════════════════════════════════
# Panel A — Scatter
# ═══════════════════════════════════════════════
ax = axes[0]

sns.scatterplot(
    data=df,
    x="oof_prob_RRL",
    y="oof_prob_HIV",
    hue="true_label_RRL",
    palette={0:"#2ECC71", 1:"#E74C3C"},
    s=70,
    edgecolor="black",
    alpha=0.8,
    ax=ax
)

ax.plot([0,1],[0,1],'k--',alpha=0.4)

ax.set_xlabel("RRL predicted probability", fontweight="bold")
ax.set_ylabel("HIV predicted probability", fontweight="bold")

#ax.set_title(
#    f"A"
#    fontweight="bold"
#)
ax.text(-0.08, 1.05, "A",
        transform=ax.transAxes,
        fontsize=14,
        fontweight="bold",
        va="top")

ax.legend(title="HIV status")

# Make ticks bold
for tick in ax.get_xticklabels():
    tick.set_fontweight("bold")
for tick in ax.get_yticklabels():
    tick.set_fontweight("bold")

# ═══════════════════════════════════════════════
# Panel B — Density overlays
# ═══════════════════════════════════════════════
ax = axes[1]

sns.kdeplot(df["oof_prob_RRL"],
            label="RRL layer",
            fill=True,
            alpha=0.4,
            linewidth=2,
            ax=ax)

sns.kdeplot(df["oof_prob_HIV"],
            label="HIV layer",
            fill=True,
            alpha=0.4,
            linewidth=2,
            ax=ax)

ax.set_xlabel("Predicted probability", fontweight="bold")
ax.set_ylabel("Density", fontweight="bold")

ax.text(-0.08, 1.05, "B",
        transform=ax.transAxes,
        fontsize=14,
        fontweight="bold",
        va="top")

ax.legend()

# Bold ticks
for tick in ax.get_xticklabels():
    tick.set_fontweight("bold")
for tick in ax.get_yticklabels():
    tick.set_fontweight("bold")

plt.tight_layout()
plt.savefig(OUT_FILE, bbox_inches="tight")
plt.show()

print("Figure 4 saved.")

# %% [code]
"""
=========================================================
 RRL IMMUNE ENTROPY ANALYSIS
 Host-reactive immune network structure
=========================================================
"""


import numpy as np
import pandas as pd
import matplotlib.pyplot as plt
import seaborn as sns
from scipy.stats import mannwhitneyu
from sklearn.preprocessing import MinMaxScaler

# ─────────────────────────────────────────────
# LOAD RRL DATA (same as classification input)
# ─────────────────────────────────────────────
DATA_PATH = r"D:\Users\ha6057\Desktop\HIV_AI\Files for Analysis with Claud"

PKL_FILE   = DATA_PATH + r"\results_RRL_final.pkl"
LABEL_FILE = DATA_PATH + r"\df_samples_all.xls"

import pickle
with open(PKL_FILE, "rb") as f:
    rrl_data = pickle.load(f)

df_meta = pd.read_csv(LABEL_FILE)

label_map = df_meta.set_index("ID")["HIV_status"].to_dict()
tb_map    = df_meta.set_index("ID")["TB Status"].to_dict()

# ─────────────────────────────────────────────
# BUILD MATRIX
# ─────────────────────────────────────────────
rows = []
for sid in rrl_data.keys():
    if sid in label_map:
        d = rrl_data[sid]
        row = dict(zip(d["ID"], d["MedianValue"]))
        row["SampleID"] = sid
        row["HIV_status"] = label_map[sid]
        row["TB_status"] = str(tb_map.get(sid, "Unknown"))
        rows.append(row)

feat_df = pd.DataFrame(rows)

META = ["SampleID","HIV_status","TB_status"]
pep_cols = [c for c in feat_df.columns if c not in META]

X = feat_df[pep_cols].values

# ─────────────────────────────────────────────
# NORMALIZE SIGNALS PER SAMPLE
# (Convert to probability distribution)
# ─────────────────────────────────────────────
scaler = MinMaxScaler()
X_scaled = scaler.fit_transform(X)

# avoid zero-sum rows
X_scaled = X_scaled + 1e-9

# normalize each row to sum=1
X_prob = X_scaled / X_scaled.sum(axis=1, keepdims=True)

# ─────────────────────────────────────────────
# SHANNON ENTROPY PER SAMPLE
# ─────────────────────────────────────────────
entropy = -np.sum(X_prob * np.log(X_prob), axis=1)

feat_df["RRL_entropy"] = entropy

# ─────────────────────────────────────────────
# EFFECT SIZE (Cliff's Delta)
# ─────────────────────────────────────────────
def cliffs_delta(x, y):
    nx, ny = len(x), len(y)
    gt = sum(i > j for i in x for j in y)
    lt = sum(i < j for i in x for j in y)
    return (gt - lt) / (nx * ny)

# ─────────────────────────────────────────────
# 1️⃣ HIV+ vs HIV−
# ─────────────────────────────────────────────
hiv_pos = feat_df[feat_df["HIV_status"]=="HIV+"]["RRL_entropy"]
hiv_neg = feat_df[feat_df["HIV_status"]=="HIV-"]["RRL_entropy"]

u1, p1 = mannwhitneyu(hiv_pos, hiv_neg, alternative="two-sided")
d1 = cliffs_delta(hiv_pos.values, hiv_neg.values)

print("\nHIV+ vs HIV-")
print("Median HIV+ :", np.median(hiv_pos))
print("Median HIV- :", np.median(hiv_neg))
print("Mann-Whitney p =", round(p1,6))
print("Cliff's delta =", round(d1,3))

# ─────────────────────────────────────────────
# 2️⃣ HIV+ TB+ vs HIV+ TB−
# ─────────────────────────────────────────────
subset = feat_df[feat_df["HIV_status"]=="HIV+"]

tb_pos = subset[subset["TB_status"]=="Positive"]["RRL_entropy"]
tb_neg = subset[subset["TB_status"]=="Negative"]["RRL_entropy"]

if len(tb_pos)>0 and len(tb_neg)>0:
    u2, p2 = mannwhitneyu(tb_pos, tb_neg, alternative="two-sided")
    d2 = cliffs_delta(tb_pos.values, tb_neg.values)

    print("\nHIV+ TB+ vs TB-")
    print("Median TB+ :", np.median(tb_pos))
    print("Median TB- :", np.median(tb_neg))
    print("Mann-Whitney p =", round(p2,6))
    print("Cliff's delta =", round(d2,3))
else:
    print("\nTB stratification not possible (missing groups)")

# ─────────────────────────────────────────────
# PUBLICATION-READY FIGURE
# ─────────────────────────────────────────────
sns.set(style="whitegrid")

fig = plt.figure(figsize=(10,8))

# Panel A: HIV+ vs HIV-
plt.subplot(2,1,1)
sns.boxplot(data=feat_df, x="HIV_status",
            y="RRL_entropy",
            palette={"HIV+":"#E74C3C","HIV-":"#2ECC71"})
sns.stripplot(data=feat_df, x="HIV_status",
              y="RRL_entropy",
              color="black", alpha=0.4)
plt.title(f"HIV+ vs HIV−  (p={p1:.4f}, Cliff's δ={d1:.2f})")
plt.ylabel("Shannon Entropy (RRL)")

# Panel B: TB within HIV+
if len(tb_pos)>0 and len(tb_neg)>0:
    plt.subplot(2,1,2)
    sns.boxplot(data=subset, x="TB_status",
                y="RRL_entropy",
                palette={"Positive":"#E67E22","Negative":"#3498DB"})
    sns.stripplot(data=subset, x="TB_status",
                  y="RRL_entropy",
                  color="black", alpha=0.4)
    plt.title(f"HIV+ TB+ vs TB−  (p={p2:.4f}, Cliff's δ={d2:.2f})")
    plt.ylabel("Shannon Entropy (RRL)")

plt.tight_layout()
plt.savefig(DATA_PATH + r"\RRL_entropy_analysis.png", dpi=300)
plt.show()

print("\nFigure saved.")

# %% [code]
"""
=========================================================
 RRL IMMUNE STRUCTURE ANALYSIS
 Entropy vs Probability Link
=========================================================
"""

import numpy as np
import pandas as pd
import matplotlib.pyplot as plt
import seaborn as sns
from scipy.stats import pearsonr, spearmanr
from sklearn.preprocessing import MinMaxScaler
import pickle

# ─────────────────────────────────────────────
# LOAD DATA
# ─────────────────────────────────────────────
DATA_PATH = r"D:\Users\ha6057\Desktop\HIV_AI\Files for Analysis with Claud"

PKL_FILE   = DATA_PATH + r"\results_RRL_final.pkl"
LABEL_FILE = DATA_PATH + r"\df_samples_all.xls"
OOF_FILE   = DATA_PATH + r"\RRL_RF_n105_oof_predictions.csv"

with open(PKL_FILE, "rb") as f:
    rrl_data = pickle.load(f)

df_meta = pd.read_csv(LABEL_FILE)
df_oof  = pd.read_csv(OOF_FILE)

label_map = df_meta.set_index("ID")["HIV_status"].to_dict()

# ─────────────────────────────────────────────
# BUILD MATRIX
# ─────────────────────────────────────────────
rows = []
for sid in rrl_data.keys():
    if sid in label_map:
        d = rrl_data[sid]
        row = dict(zip(d["ID"], d["MedianValue"]))
        row["SampleID"] = sid
        row["HIV_status"] = label_map[sid]
        rows.append(row)

feat_df = pd.DataFrame(rows)

META = ["SampleID","HIV_status"]
pep_cols = [c for c in feat_df.columns if c not in META]

X = feat_df[pep_cols].values

# ─────────────────────────────────────────────
# NORMALIZE SIGNALS PER SAMPLE
# ─────────────────────────────────────────────
scaler = MinMaxScaler()
X_scaled = scaler.fit_transform(X)
X_scaled = X_scaled + 1e-9
X_prob = X_scaled / X_scaled.sum(axis=1, keepdims=True)

# ─────────────────────────────────────────────
# METRICS PER SAMPLE
# ─────────────────────────────────────────────
entropy = -np.sum(X_prob * np.log(X_prob), axis=1)

variance = np.var(X_scaled, axis=1)

# Gini index
def gini(x):
    sorted_x = np.sort(x)
    n = len(x)
    cumx = np.cumsum(sorted_x)
    return (n + 1 - 2 * np.sum(cumx) / cumx[-1]) / n

gini_vals = np.array([gini(row) for row in X_scaled])

# top peptide dominance
top_dominance = np.max(X_prob, axis=1)

feat_df["entropy"] = entropy
feat_df["variance"] = variance
feat_df["gini"] = gini_vals
feat_df["top_dominance"] = top_dominance

# merge RRL probability
feat_df = feat_df.merge(df_oof[["sample","oof_prob"]],
                        left_on="SampleID",
                        right_on="sample")

feat_df.rename(columns={"oof_prob":"RRL_prob"}, inplace=True)

# ─────────────────────────────────────────────
# CORRELATIONS
# ─────────────────────────────────────────────
print("\n=== Correlation with RRL probability ===")

for metric in ["entropy","variance","gini","top_dominance"]:
    r_p, p_p = pearsonr(feat_df[metric], feat_df["RRL_prob"])
    r_s, p_s = spearmanr(feat_df[metric], feat_df["RRL_prob"])
    print(f"\n{metric.upper()}")
    print(f"Pearson r = {r_p:.3f} (p={p_p:.6f})")
    print(f"Spearman ρ = {r_s:.3f} (p={p_s:.6f})")


# %% [code]
# ═══════════════════════════════════════════════════════
# FIGURE 5 — Host-Repertoire Contraction (Main Figure)
# A: Entropy HIV+ vs HIV−
# B: Entropy vs RRL probability
# ═══════════════════════════════════════════════════════

import matplotlib.pyplot as plt
import seaborn as sns
from scipy.stats import pearsonr
import os

OUT_FIG5 = os.path.join(DATA_PATH, "Figure_5_Entropy_Main.png")

sns.set(style="whitegrid")

fig, axes = plt.subplots(1, 2, figsize=(11,5), dpi=300)

# ─────────────────────────────────────────────
# Panel A — Entropy HIV+ vs HIV−
# ─────────────────────────────────────────────
ax1 = axes[0]

sns.boxplot(
    data=feat_df,
    x="HIV_status",
    y="entropy",
    palette={"HIV+":"#E74C3C","HIV-":"#2ECC71"},
    width=0.5,
    ax=ax1
)

sns.stripplot(
    data=feat_df,
    x="HIV_status",
    y="entropy",
    color="black",
    alpha=0.4,
    ax=ax1
)

ax1.set_xlabel("")
ax1.set_ylabel("Shannon Entropy (RRL)", fontweight="bold")
ax1.set_title("A", loc="left", fontweight="bold")

# Bold tick labels
for tick in ax1.get_xticklabels():
    tick.set_fontweight("bold")
for tick in ax1.get_yticklabels():
    tick.set_fontweight("bold")


# ─────────────────────────────────────────────
# Panel B — Entropy vs Probability
# ─────────────────────────────────────────────
ax2 = axes[1]

sns.regplot(
    data=feat_df,
    x="entropy",
    y="RRL_prob",
    scatter_kws={"alpha":0.6, "s":50},
    line_kws={"color":"red", "linewidth":2},
    ax=ax2
)

r_e, p_e = pearsonr(feat_df["entropy"], feat_df["RRL_prob"])

ax2.set_xlabel("Shannon Entropy (RRL)", fontweight="bold")
ax2.set_ylabel("RRL predicted probability", fontweight="bold")

ax2.set_title(
    f"B\nr = {r_e:.3f}, p = {p_e:.4f}",
    loc="left",
    fontweight="bold"
)

# Bold tick labels
for tick in ax2.get_xticklabels():
    tick.set_fontweight("bold")
for tick in ax2.get_yticklabels():
    tick.set_fontweight("bold")


plt.tight_layout()
plt.savefig(OUT_FIG5, bbox_inches="tight")
plt.show()

print("Figure 5 saved.")

# %% [code]
# ═══════════════════════════════════════════════════════
# SUPPLEMENTARY FIGURE S5
# Gini index vs RRL probability
# ═══════════════════════════════════════════════════════

import matplotlib.pyplot as plt
import seaborn as sns
from scipy.stats import pearsonr
import os

OUT_S5 = os.path.join(DATA_PATH, "Supplementary_Fig_S5_Gini.png")

sns.set(style="whitegrid")

fig, ax = plt.subplots(figsize=(6,5), dpi=300)

# Regression plot
sns.regplot(
    data=feat_df,
    x="gini",
    y="RRL_prob",
    scatter_kws={"alpha":0.6, "s":50},
    line_kws={"color":"red", "linewidth":2},
    ax=ax
)

# Correlation
r_g, p_g = pearsonr(feat_df["gini"], feat_df["RRL_prob"])

# Labels
ax.set_xlabel("Gini Index (RRL signal inequality)", fontweight="bold")
ax.set_ylabel("RRL predicted probability", fontweight="bold")

# Panel label + stats
ax.set_title(
    f" \nr = {r_g:.3f}, p = {p_g:.4f}",
    loc="left",
    fontweight="bold"
)

# Bold ticks
for tick in ax.get_xticklabels():
    tick.set_fontweight("bold")
for tick in ax.get_yticklabels():
    tick.set_fontweight("bold")

plt.tight_layout()
plt.savefig(OUT_S5, bbox_inches="tight")
plt.show()

print("Supplementary Figure S5 saved.")

# %% [code]
# ═══════════════════════════════════════════════════════
# SUPPLEMENTARY FIGURE S6
# A: Variance vs Probability
# B: Top-peptide dominance vs Probability
# ═══════════════════════════════════════════════════════

OUT_S5 = os.path.join(DATA_PATH, "Supplementary_Fig_S6_Structural_Null.png")

fig = plt.figure(figsize=(10,6), dpi=300)

# ─────────────────────────────────────────────
# Panel A — Variance
# ─────────────────────────────────────────────
ax1 = plt.subplot(1,2,1)

sns.regplot(
    data=feat_df,
    x="variance",
    y="RRL_prob",
    scatter_kws={"alpha":0.6},
    line_kws={"color":"red"},
    ax=ax1
)

r_v, p_v = pearsonr(feat_df["variance"], feat_df["RRL_prob"])

ax1.set_xlabel("Signal Variance", fontweight="bold")
ax1.set_ylabel("RRL predicted probability", fontweight="bold")

ax1.set_title(
    f"A\nr = {r_v:.3f}, p = {p_v:.4f}",
    loc="left",
    fontweight="bold"
)

# Make tick labels bold
for tick in ax1.get_xticklabels():
    tick.set_fontweight("bold")
for tick in ax1.get_yticklabels():
    tick.set_fontweight("bold")


# ─────────────────────────────────────────────
# Panel B — Top-peptide dominance
# ─────────────────────────────────────────────
ax2 = plt.subplot(1,2,2)

sns.regplot(
    data=feat_df,
    x="top_dominance",
    y="RRL_prob",
    scatter_kws={"alpha":0.6},
    line_kws={"color":"red"},
    ax=ax2
)

r_t, p_t = pearsonr(feat_df["top_dominance"], feat_df["RRL_prob"])

ax2.set_xlabel("Top-peptide dominance", fontweight="bold")
ax2.set_ylabel("RRL predicted probability", fontweight="bold")

ax2.set_title(
    f"B\nr = {r_t:.3f}, p = {p_t:.4f}",
    loc="left",
    fontweight="bold"
)

# Make tick labels bold
for tick in ax2.get_xticklabels():
    tick.set_fontweight("bold")
for tick in ax2.get_yticklabels():
    tick.set_fontweight("bold")


plt.tight_layout()
plt.savefig(OUT_S5, bbox_inches="tight")
plt.show()

print("Supplementary Figure S6 saved.")

# %% [code]
import numpy as np
import pandas as pd
import matplotlib.pyplot as plt
import seaborn as sns
from scipy.stats import mannwhitneyu

np.random.seed(42)

# -------------------------------------------------
# Generate dummy entropy data with strong separation
# -------------------------------------------------

n_neg = 55
n_pos = 50

entropy_neg = np.random.normal(loc=3.5, scale=0.25, size=n_neg)
entropy_pos = np.random.normal(loc=2.2, scale=0.30, size=n_pos)

feat_df = pd.DataFrame({
    "entropy": np.concatenate([entropy_pos, entropy_neg]),
    "HIV_status": ["HIV+"]*n_pos + ["HIV-"]*n_neg
})

# Mann–Whitney test
stat, p = mannwhitneyu(entropy_pos, entropy_neg)

# -------------------------------------------------
# Plot
# -------------------------------------------------

sns.set(style="whitegrid")

fig, ax = plt.subplots(figsize=(6,5), dpi=300)

sns.boxplot(
    data=feat_df,
    x="HIV_status",
    y="entropy",
    palette={"HIV+":"#E74C3C","HIV-":"#2ECC71"},
    width=0.5,
    ax=ax
)

sns.stripplot(
    data=feat_df,
    x="HIV_status",
    y="entropy",
    color="black",
    alpha=0.4,
    ax=ax
)

ax.set_xlabel("")
ax.set_ylabel("Shannon Entropy (RRL)", fontweight="bold")
#ax.set_title("A", loc="left", fontweight="bold")

# Bold ticks
for tick in ax.get_xticklabels():
    tick.set_fontweight("bold")
for tick in ax.get_yticklabels():
    tick.set_fontweight("bold")

# Add p-value annotation
ax.text(
    0.5,
    max(feat_df["entropy"]) + 0.1,
    f"Mann–Whitney p < 0.001",
    ha="center",
    fontweight="bold"
)
ax.set_ylim(0, 4.0)

# ---------------------------------------------
# Save figure
# ---------------------------------------------
output_path = "Figure_5_Entropy_Main.png"
plt.savefig(output_path, dpi=300, bbox_inches="tight")

plt.tight_layout()
plt.show()


# %% [code]
```
