## Supplementary material for "Dual-Layer Serological Encoding: Integrating Pathogen-Derived and Host-Reactive Peptide Signatures Reveals Complementary Dimensions of HIV Immune Response": Code and Data: HIV_RF.ipynb

```
# %% [code]
"""
=============================================================================
 NESTED CROSS-VALIDATION — RANDOM FOREST + FEATURE REDUCTION
 HIV Peptide Library  |  n=105 samples  (df_samples_all.xls)
=============================================================================
 Sources:
   results_HIV_final.pkl            – peptide microarray signals (49 peptides)
   df_samples_all.xls               – HIV+/HIV- labels + metadata (n=105)
   ListeSequenses_RRL_TBC_Test.xlsx – peptide biological annotations

 Pipeline (identical methodology to n=96 analysis):
   Outer : 5-fold Stratified CV
   Inner : 5-fold Stratified CV  (k selection INSIDE training fold only)
   Model : RandomForestClassifier(n_estimators=500, class_weight='balanced')
   Feat  : Gini importance — ranked INSIDE each outer fold (no leakage)
   k grid: [3, 5, 8, 10, 15, 20, 30, 49]
   Tie   : smallest k within 1% AUC of best
   Also  : RFECV + permutation test (100 perms) + subgroup analysis
=============================================================================
"""

import pickle, warnings
import numpy  as np
import pandas as pd
import matplotlib
# matplotlib.use("Agg")  # removed — Windows uses its own backend automatically
import matplotlib.pyplot as plt
import matplotlib.patches as mpatches
from matplotlib.gridspec import GridSpec
import seaborn as sns
from collections import Counter

from sklearn.ensemble        import RandomForestClassifier
from sklearn.preprocessing   import StandardScaler
from sklearn.model_selection import (StratifiedKFold, permutation_test_score)
from sklearn.metrics         import (roc_auc_score, roc_curve,
                                     confusion_matrix,
                                     average_precision_score,
                                     precision_recall_curve)
from sklearn.feature_selection import RFECV

import warnings
from sklearn.exceptions import (
    UndefinedMetricWarning,
    FitFailedWarning,
    ConvergenceWarning
)

warnings.filterwarnings("ignore", category=UndefinedMetricWarning)
warnings.filterwarnings("ignore", category=FitFailedWarning)
warnings.filterwarnings("ignore", category=ConvergenceWarning)
warnings.filterwarnings("ignore", message="X does not have valid feature names")
warnings.filterwarnings("ignore", message="Tight layout not applied")

# ⭐ THIS is the missing line for your endless warning:
warnings.filterwarnings(
    "ignore",
    message=".*sklearn.utils.parallel.delayed.*"
)


import warnings
from sklearn.utils._testing import ignore_warnings

warnings.filterwarnings("ignore", category=UserWarning)

np.random.seed(42)
RS = 42


# ─────────────────────────────────────────────────────────────────────────────
# PATHS  ← only edit this block
# ─────────────────────────────────────────────────────────────────────────────
import os

DATA_PATH  = r"D:\Users\ha6057\Desktop\HIV_AI\Files for Analysis with Claud"
OUTPUT_PATH = DATA_PATH   # outputs saved to same folder; change if you prefer elsewhere

PKL_FILE   = os.path.join(DATA_PATH, "results_HIV_final.pkl")
LABEL_FILE = os.path.join(DATA_PATH, "df_samples_all.xls")
SEQ_FILE   = os.path.join(DATA_PATH, "ListeSequenses_RRL_TBC_Test.xlsx")

# Output files
OUT_MAIN_FIG    = os.path.join(OUTPUT_PATH, "HIV_RF_n105_Main.png")
OUT_STABLE_FIG  = os.path.join(OUTPUT_PATH, "HIV_RF_n105_StablePanel.png")
OUT_PEP_CSV     = os.path.join(OUTPUT_PATH, "HIV_RF_n105_peptides_ranked.csv")
OUT_K_CSV       = os.path.join(OUTPUT_PATH, "HIV_RF_n105_k_distribution.csv")
OUT_OOF_CSV     = os.path.join(OUTPUT_PATH, "HIV_RF_n105_oof_predictions.csv")
OUT_SG_CSV      = os.path.join(OUTPUT_PATH, "HIV_RF_n105_subgroup.csv")
# ─────────────────────────────────────────────────────────────────────────────

# ─────────────────────────────────────────────────────────────────────────────
# 1.  LOAD & MERGE
# ─────────────────────────────────────────────────────────────────────────────
print("="*70)
print("  NESTED CV  —  RF FEATURE REDUCTION  —  HIV PEPTIDE PANEL  (n=105)")
print("="*70)

with open(PKL_FILE, "rb") as f:
    hiv_data = pickle.load(f)

df_meta = pd.read_csv(LABEL_FILE)
df_meta["Country"] = df_meta["Country"].str.strip().replace({"VietNam": "Viet Nam"})

label_map   = (df_meta[df_meta["HIV_status"].isin(["HIV+","HIV-"])]
               .set_index("ID")["HIV_status"].to_dict())
tb_map      = df_meta.set_index("ID")["TB Status"].to_dict()
country_map = df_meta.set_index("ID")["Country"].to_dict()

seq_df  = pd.read_excel(SEQ_FILE)
ann_map = {}
for _, row in seq_df[seq_df["Definition"].str.contains("HIV", na=False)].iterrows():
    defn    = str(row["Definition"])
    protein = "gp41 env" if "gp41" in defn else "p24 Gag"
    ann_map[row["Sequence"]] = {
        "Name": row["Name"], "Definition": defn,
        "Protein": protein,
        "Start": row["Start"], "Stop": row["Stop"]
    }

overlap = sorted(set(hiv_data.keys()) & set(label_map.keys()))
rows = []
for sid in overlap:
    d   = hiv_data[sid]
    row = dict(zip(d["ID"], d["MedianValue"]))
    row["SampleID"]  = sid
    row["label"]     = label_map[sid]
    row["TB_status"] = str(tb_map.get(sid, "Unknown"))
    row["Country"]   = str(country_map.get(sid, "Unknown"))
    rows.append(row)

feat_df  = pd.DataFrame(rows)
META     = ["SampleID","label","TB_status","Country"]
pep_cols = [c for c in feat_df.columns if c not in META]

X = feat_df[pep_cols].values.astype(np.float64)
y = (feat_df["label"] == "HIV+").astype(int).values
n, p = X.shape

print(f"\n  Samples  : {n}  (HIV+={y.sum()}, HIV-={(y==0).sum()})")
print(f"  Peptides : {p}")
print(f"  gp41 env : {sum(1 for q in pep_cols if ann_map.get(q,{}).get('Protein','')=='gp41 env')}")
print(f"  p24 Gag  : {sum(1 for q in pep_cols if ann_map.get(q,{}).get('Protein','')=='p24 Gag')}")
print(f"  Countries: {feat_df['Country'].value_counts().to_dict()}")

# ─────────────────────────────────────────────────────────────────────────────
# 2.  NESTED 5×5 CV
# ─────────────────────────────────────────────────────────────────────────────
K_VALUES = [3, 5, 8, 10, 15, 20, 30, 49]
TIE_TOL  = 0.01

outer_cv = StratifiedKFold(n_splits=5, shuffle=True, random_state=RS)
inner_cv = StratifiedKFold(n_splits=5, shuffle=True, random_state=RS)

outer_probs      = np.zeros(n)
fold_k_opt       = []
fold_pep_idx     = []
fold_inner_aucs  = []
fold_importances = []

print(f"\n  Running nested 5x5 CV ...")
for fold_i, (tr_idx, te_idx) in enumerate(outer_cv.split(X, y)):

    X_tr, X_te = X[tr_idx], X[te_idx]
    y_tr, y_te = y[tr_idx], y[te_idx]

    # Scale on training only
    sc     = StandardScaler()
    Xtr_sc = sc.fit_transform(X_tr)
    Xte_sc = sc.transform(X_te)

    # Train RF on training set → rank peptides by Gini (NO leakage)
    rf_rank = RandomForestClassifier(
        n_estimators=500, class_weight="balanced",
        max_features="sqrt", min_samples_leaf=2,
        random_state=RS, n_jobs=1)
    rf_rank.fit(Xtr_sc, y_tr)
    imp  = rf_rank.feature_importances_
    rank = np.argsort(-imp)
    fold_importances.append(imp.copy())

    # Inner CV: evaluate each k strictly on training data
    k_auc = {}
    for k in K_VALUES:
        Xtr_k  = Xtr_sc[:, rank[:k]]
        scores = []
        for in_tr, in_val in inner_cv.split(Xtr_k, y_tr):
            if len(np.unique(y_tr[in_val])) < 2:
                continue
            rf_in = RandomForestClassifier(
                n_estimators=300, class_weight="balanced",
                max_features="sqrt", min_samples_leaf=2,
                random_state=RS, n_jobs=1)
            rf_in.fit(Xtr_k[in_tr], y_tr[in_tr])
            pv = rf_in.predict_proba(Xtr_k[in_val])[:, 1]
            if len(np.unique(y_tr[in_val])) >= 2:
                scores.append(roc_auc_score(y_tr[in_val], pv))
        k_auc[k] = np.mean(scores) if scores else 0.0
    fold_inner_aucs.append(k_auc)

    # Select k_opt: best inner AUC, tie-break to smallest k
    best_v = max(k_auc.values())
    k_opt  = min([k for k, v in k_auc.items() if v >= best_v - TIE_TOL])
    fold_k_opt.append(k_opt)
    fold_pep_idx.append(rank[:k_opt].tolist())

    # Retrain with k_opt, predict test fold
    top_opt = rank[:k_opt]
    rf_f = RandomForestClassifier(
        n_estimators=500, class_weight="balanced",
        max_features="sqrt", min_samples_leaf=2,
        random_state=RS, n_jobs=1)
    rf_f.fit(Xtr_sc[:, top_opt], y_tr)
    outer_probs[te_idx] = rf_f.predict_proba(Xte_sc[:, top_opt])[:, 1]

    fold_auc = roc_auc_score(y_te, outer_probs[te_idx])
    print(f"    Fold {fold_i+1}/5 | k_opt={k_opt:2d} | "
          f"inner_AUC={best_v:.3f} | outer_AUC={fold_auc:.3f}")

# ─────────────────────────────────────────────────────────────────────────────
# 3.  AGGREGATE PERFORMANCE
# ─────────────────────────────────────────────────────────────────────────────
final_auc      = roc_auc_score(y, outer_probs)
ap_score       = average_precision_score(y, outer_probs)
fpr, tpr, thrs = roc_curve(y, outer_probs)
j              = np.argmax(tpr - fpr)
opt_thr        = thrs[j]
y_pred         = (outer_probs >= opt_thr).astype(int)
tn, fp, fn, tp_v = confusion_matrix(y, y_pred).ravel()
sens = tp_v / (tp_v + fn)
spec = tn   / (tn   + fp)
ppv  = tp_v / (tp_v + fp) if tp_v + fp else 0
npv  = tn   / (tn   + fn) if tn   + fn else 0
f1   = 2 * tp_v / (2 * tp_v + fp + fn)
mcc_n = tp_v * tn - fp * fn
mcc_d = np.sqrt((tp_v+fp)*(tp_v+fn)*(tn+fp)*(tn+fn))
mcc   = mcc_n / mcc_d if mcc_d else 0

k_counts       = Counter(fold_k_opt)
best_k         = k_counts.most_common(1)[0][0]
mean_inner_auc = {k: np.mean([d[k] for d in fold_inner_aucs]) for k in K_VALUES}
std_inner_auc  = {k: np.std( [d[k] for d in fold_inner_aucs]) for k in K_VALUES}
mean_imp       = np.mean(fold_importances, axis=0)
std_imp        = np.std( fold_importances, axis=0)
global_rank    = np.argsort(-mean_imp)

# Per-fold outer AUCs for plotting
fold_outer_aucs = []
for fold_i, (_, te_idx) in enumerate(outer_cv.split(X, y)):
    if len(np.unique(y[te_idx])) == 2:
        fold_outer_aucs.append(roc_auc_score(y[te_idx], outer_probs[te_idx]))
    else:
        fold_outer_aucs.append(np.nan)

# ─────────────────────────────────────────────────────────────────────────────
# 4.  STABLE PEPTIDES  (>=60% selection frequency)
# ─────────────────────────────────────────────────────────────────────────────
freq_cnt  = Counter()
for idx_l in fold_pep_idx:
    freq_cnt.update(idx_l)

# number of outer folds
n_folds = outer_cv.get_n_splits()

# stability rule: selected in ≥3 out of 5 folds
min_folds = 3
stable_thr = 100.0 * min_folds / n_folds

stable_peps = sorted(
    [pep_cols[i] for i, c in freq_cnt.items() if c >= min_folds],
    key=lambda pep: -mean_imp[pep_cols.index(pep)]
)

# compute percentage only for reporting (not selection)
freq_pct = {pep_cols[i]: 100.0 * c / n_folds for i, c in freq_cnt.items()}

# ─────────────────────────────────────────────────────────────────────────────
# 5.  RFECV
# ─────────────────────────────────────────────────────────────────────────────
print("\n  Running RFECV ...")
sc_rfecv = StandardScaler()
X_sc     = sc_rfecv.fit_transform(X)
rfecv    = RFECV(
    RandomForestClassifier(n_estimators=200, class_weight="balanced",
                            min_samples_leaf=2, random_state=RS, n_jobs=1),
    step=1,
    cv=StratifiedKFold(5, shuffle=True, random_state=RS),
    scoring="roc_auc",
    min_features_to_select=3, n_jobs=1)
rfecv.fit(X_sc, y)
rfecv_n    = rfecv.n_features_
rfecv_peps = [pep_cols[i] for i in range(p) if rfecv.support_[i]]
print(f"  RFECV optimal: {rfecv_n} peptides | "
      f"AUC={rfecv.cv_results_['mean_test_score'].max():.4f}")

# ─────────────────────────────────────────────────────────────────────────────
# 6.  FINAL MODEL on stable peptides
# ─────────────────────────────────────────────────────────────────────────────
stable_idx  = [pep_cols.index(pep) for pep in stable_peps]
X_stable    = X[:, stable_idx]
sc_f        = StandardScaler()
X_stable_sc = sc_f.fit_transform(X_stable)
rf_final    = RandomForestClassifier(
    n_estimators=1000, class_weight="balanced",
    max_features="sqrt", min_samples_leaf=2,
    random_state=RS, n_jobs=-1)
rf_final.fit(X_stable_sc, y)
final_imp_stable = dict(zip(stable_peps, rf_final.feature_importances_))
probs_full       = rf_final.predict_proba(X_stable_sc)[:, 1]

# ─────────────────────────────────────────────────────────────────────────────
# 7.  SUBGROUP ANALYSIS
# ─────────────────────────────────────────────────────────────────────────────
sg_rows = []
for col in ["TB_status", "Country"]:
    for val in feat_df[col].unique():
        mask = (feat_df[col] == val).values
        if mask.sum() >= 6 and len(np.unique(y[mask])) == 2:
            sg_rows.append({
                "Group":   f"{col}={val}",
                "N":       mask.sum(),
                "HIV+":    int(y[mask].sum()),
                "AUC":     round(roc_auc_score(y[mask], probs_full[mask]), 3)
            })
sg_df = pd.DataFrame(sg_rows).sort_values("AUC", ascending=False)

# ─────────────────────────────────────────────────────────────────────────────
# 8.  PERMUTATION TEST
# ─────────────────────────────────────────────────────────────────────────────
print("  Running permutation test (100 permutations) ...")
obs_auc, perm_aucs, pval = permutation_test_score(
    RandomForestClassifier(n_estimators=200, class_weight="balanced",
                            min_samples_leaf=2, random_state=RS, n_jobs=1),
    X_stable_sc, y,
    cv=StratifiedKFold(5, shuffle=True, random_state=RS),
    scoring="roc_auc", n_permutations=100,
    random_state=RS, n_jobs=1)
print(f"  Observed AUC = {obs_auc:.4f}  |  p = {pval:.4f}")

# ─────────────────────────────────────────────────────────────────────────────
# 9.  CONSOLE REPORT
# ─────────────────────────────────────────────────────────────────────────────
print("\n" + "="*70)
print("  RESULTS")
print("="*70)

print(f"""
  +------------------------------------------------------------------+
  |  NESTED 5x5-FOLD CV PERFORMANCE  (leakage-free, n=105)          |
  |                                                                  |
  |  AUC (pooled OOF)   = {final_auc:.4f}                               |
  |  Avg Precision      = {ap_score:.4f}                               |
  |  Sensitivity        = {sens:.4f}  ({tp_v} TP / {tp_v+fn} HIV+)              |
  |  Specificity        = {spec:.4f}  ({tn} TN / {tn+fp} HIV-)              |
  |  PPV                = {ppv:.4f}                               |
  |  NPV                = {npv:.4f}                               |
  |  F1                 = {f1:.4f}                               |
  |  MCC                = {mcc:.4f}                               |
  |  Threshold (Youden) = {opt_thr:.4f}                               |
  +------------------------------------------------------------------+""")

print(f"\n  Optimal k per outer fold:")
for i, (k, ki) in enumerate(zip(fold_k_opt, fold_inner_aucs)):
    print(f"    Fold {i+1}: k={k:2d}  inner_AUC={ki[k]:.3f}  outer_AUC={fold_outer_aucs[i]:.3f}")

print(f"\n  Inner CV AUC by k (mean +/- SD over 5 folds):")
for k in K_VALUES:
    mark = "  <- selected" if k == best_k else ""
    print(f"    k={k:2d}  {mean_inner_auc[k]:.4f} +/- {std_inner_auc[k]:.4f}{mark}")

print(f"\n  RFECV optimal: {rfecv_n} peptides")

print(f"\n  Stable peptides (>={stable_thr:.0f}% selection, n={len(stable_peps)}):")
print(f"    {'#':>3}  {'Peptide':<22}  {'Protein':<12}  {'Freq':>6}  {'MeanGini':>9}  {'RFECV':>6}")
for i, pep in enumerate(stable_peps, 1):
    prot = ann_map.get(pep, {}).get("Protein", "?")
    fr   = freq_pct.get(pep, 0)
    mi   = mean_imp[pep_cols.index(pep)]
    in_r = "yes" if pep in rfecv_peps else "no"
    print(f"    {i:>3}  {pep:<22}  {prot:<12}  {fr:>5.0f}%  {mi:.6f}  {in_r:>6}")

print(f"\n  Subgroup AUC (stable peptide model):")
print(sg_df.to_string(index=False))
print(f"\n  Permutation test: p = {pval:.4f}  (observed AUC = {obs_auc:.4f})")

# ─────────────────────────────────────────────────────────────────────────────
# 10.  FIGURES
# ─────────────────────────────────────────────────────────────────────────────
C = dict(pos="#E74C3C", neg="#2ECC71", main="#2C3E50",
         acc="#E67E22", blue="#3498DB", gp41="#8E44AD",
         p24="#2980B9", bg="#F8F9FA", gray="#95A5A6")

def prot_color(pep):
    return C["gp41"] if ann_map.get(pep, {}).get("Protein","") == "gp41 env" else C["p24"]

# ══════════════════════════════════════════════════════════════════════════════
#  FIGURE 1 — Main results dashboard
# ══════════════════════════════════════════════════════════════════════════════
fig1 = plt.figure(figsize=(22, 26))
fig1.patch.set_facecolor(C["bg"])
gs1  = GridSpec(4, 3, figure=fig1, hspace=0.46, wspace=0.36,
                left=0.07, right=0.97, top=0.94, bottom=0.04)

# A  ROC curve
ax = fig1.add_subplot(gs1[0, 0])
ax.plot(fpr, tpr, lw=3, color=C["main"],
        label=f"Random Forest  AUC = {final_auc:.4f}")
ax.fill_between(fpr, tpr, alpha=0.12, color=C["main"])
ax.plot([0,1],[0,1], "k--", lw=1.2, alpha=0.45, label="Random")
ax.scatter([1-spec], [sens], s=150, zorder=6, color=C["pos"],
           label=f"Op.pt  Sens={sens:.3f}, Spec={spec:.3f}")
ax.annotate(f"Thr={opt_thr:.3f}", xy=(1-spec, sens),
            xytext=(1-spec+0.13, sens-0.11),
            arrowprops=dict(arrowstyle="->", color=C["pos"], lw=1.3),
            fontsize=9, color=C["pos"])
ax.set(xlabel="1 - Specificity (FPR)", ylabel="Sensitivity (TPR)",
       title="A   ROC Curve\n(nested 5-fold CV, leakage-free, n=105)",
       xlim=(-0.02,1.02), ylim=(-0.02,1.05))
ax.legend(fontsize=8.5, loc="lower right")
ax.grid(alpha=0.3); ax.set_facecolor("white")

# B  Precision-Recall
ax = fig1.add_subplot(gs1[0, 1])
prec, rec, _ = precision_recall_curve(y, outer_probs)
ax.plot(rec, prec, lw=2.5, color=C["acc"],
        label=f"RF  AP = {ap_score:.4f}")
ax.axhline(y.mean(), color="gray", lw=1.5, ls="--",
           label=f"Baseline = {y.mean():.2f}")
ax.fill_between(rec, prec, alpha=0.12, color=C["acc"])
ax.set(xlabel="Recall", ylabel="Precision",
       title="B   Precision-Recall Curve",
       xlim=(-0.02,1.02), ylim=(-0.02,1.05))
ax.legend(fontsize=9); ax.grid(alpha=0.3); ax.set_facecolor("white")

# C  AUC vs k
ax = fig1.add_subplot(gs1[0, 2])
ks   = K_VALUES
aucs = [mean_inner_auc[k] for k in ks]
stds = [std_inner_auc[k]  for k in ks]
ax.plot(ks, aucs, "o-", color=C["main"], lw=2.5, ms=9, zorder=5)
ax.fill_between(ks,
                [a-s for a,s in zip(aucs, stds)],
                [a+s for a,s in zip(aucs, stds)],
                alpha=0.18, color=C["main"], label="Mean +/- 1 SD")
ax.axvline(best_k, color=C["acc"], lw=2.2, ls="--",
           label=f"Optimal k = {best_k}")
ax.scatter([best_k], [mean_inner_auc[best_k]], s=160, color=C["acc"], zorder=7)
for k, a in zip(ks, aucs):
    ax.annotate(f"{a:.3f}", xy=(k, a), xytext=(0, 9),
                textcoords="offset points", ha="center",
                fontsize=8.5, color=C["main"])
ax.set(xlabel="k (peptides)", ylabel="Mean inner-CV AUC",
       title="C   AUC vs k\n(inner CV, 5 outer folds)",
       xticks=ks, ylim=(0.75, 1.02))
ax.legend(fontsize=9); ax.grid(alpha=0.3); ax.set_facecolor("white")

# D  Feature importance top-20
ax = fig1.add_subplot(gs1[1, 0])
top20_idx  = global_rank[:20]
top20_imp  = mean_imp[top20_idx][::-1]
top20_std  = std_imp[top20_idx][::-1]
top20_peps = [pep_cols[i] for i in top20_idx][::-1]
top20_cols = [prot_color(q) for q in top20_peps]
ax.barh(range(20), top20_imp, xerr=top20_std,
        color=top20_cols, edgecolor="white",
        height=0.75, alpha=0.88, capsize=3)
ax.set_yticks(range(20))
ax.set_yticklabels(top20_peps, fontsize=8.5, fontfamily="monospace")
ax.set(xlabel="Mean Gini importance (+/- SD)",
       title="D   Top-20 Peptides by Importance\n(consensus across 5 outer folds)")
gp41_p = mpatches.Patch(color=C["gp41"], label="gp41 env")
p24_p  = mpatches.Patch(color=C["p24"],  label="p24 Gag")
ax.legend(handles=[gp41_p, p24_p], fontsize=9)
ax.grid(axis="x", alpha=0.3); ax.set_facecolor("white")

# E  Confusion matrix
ax = fig1.add_subplot(gs1[1, 1])
cm_arr = np.array([[tn, fp],[fn, tp_v]])
sns.heatmap(cm_arr, annot=True, fmt="d", cmap="Greens",
            xticklabels=["Pred HIV-","Pred HIV+"],
            yticklabels=["Actual HIV-","Actual HIV+"],
            ax=ax, cbar=False, linewidths=1.5, linecolor="white",
            annot_kws={"size":18,"weight":"bold"})
ax.set_title("E   Confusion Matrix\n(nested CV, Youden threshold)",
             fontsize=12, fontweight="bold")
mt = (f"Sens = {sens:.3f}\nSpec = {spec:.3f}\n"
      f"PPV  = {ppv:.3f}\nNPV  = {npv:.3f}\n"
      f"F1   = {f1:.3f}\nMCC  = {mcc:.3f}")
ax.text(1.08, 0.5, mt, transform=ax.transAxes, fontsize=11, va="center",
        bbox=dict(boxstyle="round", facecolor="#EAFAF1", alpha=0.9))

# F  Score distribution
ax = fig1.add_subplot(gs1[1, 2])
bins = np.linspace(0, 1, 26)
ax.hist(outer_probs[y==0], bins, alpha=0.65, color=C["neg"],
        edgecolor="white", label=f"HIV-  (n={(y==0).sum()})")
ax.hist(outer_probs[y==1], bins, alpha=0.65, color=C["pos"],
        edgecolor="white", label=f"HIV+  (n={y.sum()})")
ax.axvline(opt_thr, color="black", lw=2, ls="--",
           label=f"Threshold = {opt_thr:.3f}")
ax.set(xlabel="Predicted probability (HIV+)", ylabel="Count",
       title="F   Score Distribution\n(out-of-fold, n=105)")
ax.legend(fontsize=9); ax.grid(alpha=0.3); ax.set_facecolor("white")

# G  Stability frequency all peptides
ax = fig1.add_subplot(gs1[2, :])
srt      = sorted(freq_pct.items(), key=lambda x: -x[1])
all_peps_s = [x[0] for x in srt]
all_freq_s = [x[1] for x in srt]
bar_c    = []
for pep, f in zip(all_peps_s, all_freq_s):
    if f >= stable_thr:
        bar_c.append(prot_color(pep))
    elif f >= 40:
        bar_c.append(C["blue"])
    else:
        bar_c.append("lightgray")
ax.bar(range(len(all_peps_s)), all_freq_s, color=bar_c,
       edgecolor="white", alpha=0.88)
ax.axhline(stable_thr, color=C["acc"], lw=2.2, ls="--",
           label=f"Stability threshold ({stable_thr:.0f}%)")
ax.set_xticks(range(len(all_peps_s)))
ax.set_xticklabels([pep[:16] for pep in all_peps_s], rotation=65, ha="right",
                    fontsize=8, fontfamily="monospace")
for i, (pep, f) in enumerate(zip(all_peps_s, all_freq_s)):
    if f >= stable_thr:
        ax.text(i, f+1.2, f"{f:.0f}%", ha="center",
                fontsize=8, color=prot_color(pep), fontweight="bold")
ax.set(ylabel="Selection frequency (%)",
       title="G   Peptide Stability — Selection Frequency Across 5 Outer Folds  (n=105)")
h_gp41 = mpatches.Patch(color=C["gp41"], label=f"gp41 env, stable >={stable_thr:.0f}%")
h_p24  = mpatches.Patch(color=C["p24"],  label=f"p24 Gag, stable >={stable_thr:.0f}%")
h_freq = mpatches.Patch(color=C["blue"], label="Frequent >=40%")
h_low  = mpatches.Patch(color="lightgray", label="<40%")
ax.legend(handles=[h_gp41, h_p24, h_freq, h_low, ax.lines[0]],
          fontsize=9, loc="upper right")
ax.grid(axis="y", alpha=0.3); ax.set_facecolor("white")

# H  RFECV curve
ax = fig1.add_subplot(gs1[3, 0])
nf_ = rfecv.cv_results_["n_features"]
ms_ = rfecv.cv_results_["mean_test_score"]
ss_ = rfecv.cv_results_["std_test_score"]
ax.plot(nf_, ms_, color=C["main"], lw=2.2)
ax.fill_between(nf_, ms_-ss_, ms_+ss_, alpha=0.15, color=C["main"])
ax.axvline(rfecv_n, color=C["acc"], lw=2, ls="--",
           label=f"RFECV optimal: {rfecv_n}")
ax.set(xlabel="Number of features", ylabel="CV AUC",
       title=f"H   RFECV Curve\n(optimal: {rfecv_n} peptides)")
ax.legend(fontsize=9); ax.grid(alpha=0.3); ax.set_facecolor("white")

# I  Permutation test
ax = fig1.add_subplot(gs1[3, 1])
ax.hist(perm_aucs, bins=20, color=C["blue"],
        alpha=0.7, edgecolor="white", label="Permuted AUCs")
ax.axvline(obs_auc, color=C["pos"], lw=2.5,
           label=f"Observed = {obs_auc:.4f}")
ax.set(xlabel="AUC", ylabel="Count",
       title=f"I   Permutation Test\n(100 perms, p = {pval:.4f})")
ax.legend(fontsize=9); ax.grid(alpha=0.3); ax.set_facecolor("white")

# J  Subgroup AUC
ax = fig1.add_subplot(gs1[3, 2])
sg_s = sg_df.sort_values("AUC", ascending=True)
bcols_sg = [C["pos"] if "TB" in g else C["blue"] for g in sg_s["Group"]]
ax.barh(range(len(sg_s)), sg_s["AUC"], color=bcols_sg,
        height=0.6, alpha=0.85, edgecolor="white")
ax.set_yticks(range(len(sg_s)))
ax.set_yticklabels(sg_s["Group"], fontsize=9)
for i, row in sg_s.reset_index(drop=True).iterrows():
    ax.text(row["AUC"]+0.005, i,
            f"n={row['N']} ({row['HIV+']}+)", va="center", fontsize=8.5)
ax.axvline(0.8, color="gray", lw=1, ls="--", alpha=0.5)
ax.set(xlabel="AUC",
       title="J   Subgroup Analysis\n(stable peptide model)",
       xlim=(0.55, 1.10))
h1 = mpatches.Patch(color=C["pos"],  label="TB subgroup")
h2 = mpatches.Patch(color=C["blue"], label="Country subgroup")
ax.legend(handles=[h1,h2], fontsize=9)
ax.grid(axis="x", alpha=0.3); ax.set_facecolor("white")

fig1.suptitle(
    f"Nested 5x5-Fold CV  —  Random Forest + Feature Reduction  —  HIV Peptide Panel\n"
    f"n=105 samples  |  {p} peptides  |  "
    f"Outer AUC = {final_auc:.4f}  |  Optimal k = {best_k}  |  "
    f"Stable panel = {len(stable_peps)} peptides",
    fontsize=14, fontweight="bold", y=0.97, color=C["main"])

fig1.savefig(OUT_MAIN_FIG,
             dpi=150, bbox_inches="tight", facecolor=C["bg"])
plt.close(fig1)
print("\n  Fig 1 saved.")

# ══════════════════════════════════════════════════════════════════════════════
#  FIGURE 2 — Stable panel deep dive
# ══════════════════════════════════════════════════════════════════════════════
fig2 = plt.figure(figsize=(22, 16))
fig2.patch.set_facecolor(C["bg"])
gs2  = GridSpec(2, 3, figure=fig2, hspace=0.44, wspace=0.36,
                left=0.07, right=0.97, top=0.92, bottom=0.07)

# A  Importance + frequency dual bar
ax  = fig2.add_subplot(gs2[0, :2])
ax2 = ax.twinx()
stab_ord = sorted(stable_peps, key=lambda pep: -mean_imp[pep_cols.index(pep)])
s_imp    = [mean_imp[pep_cols.index(pep)] for pep in stab_ord]
s_std    = [std_imp[pep_cols.index(pep)]  for pep in stab_ord]
s_freq   = [freq_pct.get(pep, 0)          for pep in stab_ord]
s_cols   = [prot_color(pep)               for pep in stab_ord]
imp_norm = np.array(s_imp) / max(s_imp)
x_pos    = np.arange(len(stab_ord))

ax.bar(x_pos - 0.2, imp_norm, 0.36, color=s_cols,
       alpha=0.85, edgecolor="white")
ax.errorbar(x_pos - 0.2, imp_norm,
            yerr=[se/max(s_imp) for se in s_std],
            fmt="none", color="black", capsize=4, lw=1.2)
ax2.bar(x_pos + 0.2, s_freq, 0.36, color=C["acc"],
        alpha=0.80, edgecolor="white")
ax.set_xticks(x_pos)
ax.set_xticklabels(stab_ord, rotation=25, ha="right",
                    fontsize=10, fontfamily="monospace")
for i, (iv, fv) in enumerate(zip(imp_norm, s_freq)):
    ax.text(i-0.2,  iv+0.02, f"{iv:.2f}", ha="center",
            fontsize=9, color="black", fontweight="bold")
    ax2.text(i+0.2, fv+1.0,  f"{fv:.0f}%", ha="center",
             fontsize=9, color=C["acc"], fontweight="bold")
ax.set_ylabel("Gini importance (normalised)", color=C["main"], fontsize=11)
ax2.set_ylabel("Selection frequency (%)",     color=C["acc"],  fontsize=11)
ax.tick_params(axis="y", labelcolor=C["main"])
ax2.tick_params(axis="y", labelcolor=C["acc"])
ax.set_title("A   Stable Peptides — Importance vs Selection Frequency  (n=105)",
             fontsize=12, fontweight="bold")
h_gp41_  = mpatches.Patch(color=C["gp41"], alpha=0.85, label="gp41 env")
h_p24_   = mpatches.Patch(color=C["p24"],  alpha=0.85, label="p24 Gag")
h_freq_  = mpatches.Patch(color=C["acc"],  alpha=0.80, label="Selection freq (%)")
ax.legend(handles=[h_gp41_, h_p24_, h_freq_], fontsize=9, loc="upper right")
ax.set_facecolor("white")

# B  Signal box plots top-6
ax = fig2.add_subplot(gs2[0, 2])
top6 = stab_ord[:6]
for i, pep in enumerate(top6):
    dp = X[:, pep_cols.index(pep)][y==1]
    dn = X[:, pep_cols.index(pep)][y==0]
    jp = np.random.normal(i-0.18, 0.05, len(dp))
    jn = np.random.normal(i+0.18, 0.05, len(dn))
    ax.scatter(jp, dp, alpha=0.45, s=22, color=C["pos"], zorder=3)
    ax.scatter(jn, dn, alpha=0.45, s=22, color=C["neg"], zorder=3)
    bp = ax.boxplot([dp], positions=[i-0.18], widths=0.28, patch_artist=True,
                    manage_ticks=False, medianprops=dict(color="white", lw=2))
    bn = ax.boxplot([dn], positions=[i+0.18], widths=0.28, patch_artist=True,
                    manage_ticks=False, medianprops=dict(color="white", lw=2))
    bp["boxes"][0].set(facecolor=C["pos"], alpha=0.55)
    bn["boxes"][0].set(facecolor=C["neg"], alpha=0.55)
ax.set_xticks(range(6))
ax.set_xticklabels([pep[:14] for pep in top6], rotation=35, ha="right",
                    fontsize=8.5, fontfamily="monospace")
ax.set(ylabel="Median Signal", title="B   Signal Distributions\n(top-6 stable peptides)")
h1 = mpatches.Patch(color=C["pos"], alpha=0.7, label="HIV+")
h2 = mpatches.Patch(color=C["neg"], alpha=0.7, label="HIV-")
ax.legend(handles=[h1,h2], fontsize=9)
ax.grid(axis="y", alpha=0.3); ax.set_facecolor("white")

# C  Per-fold outer AUC
ax = fig2.add_subplot(gs2[1, 0])
fold_nums = [f"Fold {i+1}" for i in range(5)]
colors_f  = [C["main"] if not np.isnan(a) else "gray" for a in fold_outer_aucs]
ax.bar(fold_nums, fold_outer_aucs, color=colors_f,
       edgecolor="white", alpha=0.85, width=0.55)
for i, (fn_, a) in enumerate(zip(fold_nums, fold_outer_aucs)):
    if not np.isnan(a):
        ax.text(i, a+0.005, f"{a:.3f}", ha="center",
                fontsize=11, fontweight="bold", color=C["main"])
ax.axhline(final_auc, color=C["acc"], lw=2, ls="--",
           label=f"Overall AUC = {final_auc:.4f}")
ax.set(ylabel="AUC", title="C   Per-Fold Outer AUC", ylim=(0.6, 1.08))
ax.legend(fontsize=9); ax.grid(axis="y", alpha=0.3); ax.set_facecolor("white")

# D  Annotation table
ax = fig2.add_subplot(gs2[1, 1:])
ax.axis("off")
table_data = []
for i, pep in enumerate(stable_peps, 1):
    a = ann_map.get(pep, {})
    table_data.append([
        str(i), pep,
        a.get("Protein", "?"),
        a.get("Name", "?"),
        f"{a.get('Start','?')}-{a.get('Stop','?')}",
        f"{freq_pct.get(pep,0):.0f}%",
        f"{mean_imp[pep_cols.index(pep)]:.5f}",
        "yes" if pep in rfecv_peps else "no"
    ])
col_labels = ["#","Sequence","Protein","Name","Position","Stability","Gini Imp","RFECV"]
tbl = ax.table(cellText=table_data, colLabels=col_labels,
               cellLoc="center", loc="center",
               bbox=[0, 0.02, 1, 0.94])
tbl.auto_set_font_size(False); tbl.set_fontsize(8.5)
for (r, col_), cell in tbl.get_celld().items():
    if r == 0:
        cell.set(facecolor=C["main"])
        cell.set_text_props(color="white", fontweight="bold")
    elif r <= len(table_data):
        prot_r = table_data[r-1][2]
        cell.set(facecolor="#F5EEF8" if "gp41" in prot_r else "#EBF5FB")
    cell.set_edgecolor("white")
ax.set_title("D   Stable Peptide Biological Annotations",
             fontsize=12, fontweight="bold", pad=15)

fig2.suptitle(
    f"Stable Panel — {len(stable_peps)} Peptides Selected in >={stable_thr:.0f}% of Outer Folds  (n=105)",
    fontsize=14, fontweight="bold", y=1.01, color=C["main"])
plt.tight_layout()
fig2.savefig(OUT_STABLE_FIG,
             dpi=150, bbox_inches="tight", facecolor=C["bg"])
plt.close(fig2)
print("  Fig 2 saved.")

# ─────────────────────────────────────────────────────────────────────────────
# 11.  CSV OUTPUTS
# ─────────────────────────────────────────────────────────────────────────────

# All peptides ranked
all_pep_rows = []
for i, idx in enumerate(global_rank):
    pep = pep_cols[idx]
    a   = ann_map.get(pep, {})
    all_pep_rows.append({
        "rank":               i+1,
        "peptide":            pep,
        "protein":            a.get("Protein","?"),
        "name":               a.get("Name","?"),
        "position":           f"{a.get('Start','?')}-{a.get('Stop','?')}",
        "mean_gini_imp":      round(mean_imp[idx], 6),
        "std_gini_imp":       round(std_imp[idx],  6),
        "selection_freq_pct": round(freq_pct.get(pep, 0), 1),
        "stable":             freq_pct.get(pep, 0) >= stable_thr,
        "rfecv_selected":     pep in rfecv_peps,
    })
pd.DataFrame(all_pep_rows).to_csv(OUT_PEP_CSV, index=False)

# k distribution
pd.DataFrame([{
    "k":                k,
    "n_folds_selected": k_counts.get(k, 0),
    "pct_folds":        round(100 * k_counts.get(k, 0) / 5, 1),
    "mean_inner_AUC":   round(mean_inner_auc[k], 4),
    "std_inner_AUC":    round(std_inner_auc[k],  4),
} for k in K_VALUES]).to_csv(OUT_K_CSV, index=False)

# OOF predictions
pd.DataFrame({
    "sample":     feat_df["SampleID"].tolist(),
    "true_label": y,
    "oof_prob":   np.round(outer_probs, 6),
    "predicted":  y_pred,
    "TB_status":  feat_df["TB_status"].tolist(),
    "Country":    feat_df["Country"].tolist(),
}).to_csv(OUT_OOF_CSV, index=False)

# Subgroup
sg_df.to_csv(OUT_SG_CSV, index=False)

print("  CSVs saved.")
print("\n  ALL DONE.")


# %% [code]
display(fig1) 

# %% [code]
display(fig2) 

# %% [code]
# ─────────────────────────────────────────────────────────────
# HEATMAP — Stable Peptides (Z-scored) — Clean Journal Version
# ─────────────────────────────────────────────────────────────

import numpy as np
import pandas as pd
import matplotlib.pyplot as plt
import seaborn as sns
from matplotlib.colors import TwoSlopeNorm

# ---------- 1. Prepare matrix (stable peptides only)

stable_idx = [pep_cols.index(p) for p in stable_peps]
X_stable   = X[:, stable_idx]

# Z-score across samples (per peptide)
X_z = (X_stable - X_stable.mean(axis=0)) / X_stable.std(axis=0)

# Clip at ±3 SD
X_z = np.clip(X_z, -3, 3)

# ---------- 2. Order samples

df_heat = pd.DataFrame(X_z, columns=stable_peps)
df_heat["HIV"]  = y
df_heat["prob"] = outer_probs

df_pos = df_heat[df_heat["HIV"] == 1].sort_values("prob", ascending=False)
df_neg = df_heat[df_heat["HIV"] == 0].sort_values("prob", ascending=False)

df_sorted = pd.concat([df_pos, df_neg])
heat_matrix = df_sorted[stable_peps].T.values

n_pos = df_pos.shape[0]
n_neg = df_neg.shape[0]

# ---------- 3. Create heatmap

plt.figure(figsize=(16, 6), dpi=150)

norm = TwoSlopeNorm(vmin=-3, vcenter=0, vmax=3)

ax = sns.heatmap(
    heat_matrix,
    cmap="RdBu_r",
    norm=norm,
    xticklabels=False,
    yticklabels=stable_peps,
    cbar_kws={"label": "Z-score (IgG signal)"}
)

# ---------- 4. Vertical separator

plt.axvline(n_pos, color="black", linewidth=2)

# ---------- 5. Make all labels bold & black

# Y-axis label
ax.set_ylabel("Peptides", fontsize=12, fontweight="bold", color="black")

# Remove X-axis label
ax.set_xlabel("")

# Peptide labels
for label in ax.get_yticklabels():
    label.set_fontweight("bold")
    label.set_color("black")

# Remove X ticks completely
ax.set_xticks([])

# Colorbar formatting
cbar = ax.collections[0].colorbar
cbar.ax.yaxis.label.set_fontweight("bold")
cbar.ax.yaxis.label.set_color("black")

for t in cbar.ax.get_yticklabels():
    t.set_fontweight("bold")
    t.set_color("black")

# ---------- 6. Remove title (do nothing)

plt.tight_layout()

# ---------- 7. Save figure

plt.savefig(
    "HIV_StablePanel_Heatmap_Clean.png",
    dpi=300,
    bbox_inches="tight"
)

plt.show()

# %% [code]
import pandas as pd
import numpy as np
import matplotlib.pyplot as plt
from matplotlib.gridspec import GridSpec
from sklearn.metrics import roc_curve, roc_auc_score, confusion_matrix
import os

# ---- PATH ----
OUTPUT_PATH = r"D:\Users\ha6057\Desktop\HIV_AI\Files for Analysis with Claud"
OOF_FILE = os.path.join(OUTPUT_PATH, "HIV_RF_n105_oof_predictions.csv")
PEP_FILE = os.path.join(OUTPUT_PATH, "HIV_RF_n105_peptides_ranked.csv")

# ---- LOAD OOF DATA ----
df_oof = pd.read_csv(OOF_FILE)

y = df_oof["true_label"].values
outer_probs = df_oof["oof_prob"].values

# ---- ROC ----
final_auc = roc_auc_score(y, outer_probs)
fpr, tpr, thresholds = roc_curve(y, outer_probs)

j = np.argmax(tpr - fpr)
opt_thr = thresholds[j]

y_pred = (outer_probs >= opt_thr).astype(int)
tn, fp, fn, tp = confusion_matrix(y, y_pred).ravel()
sens = tp / (tp + fn)
spec = tn / (tn + fp)

# ---- LOAD PEPTIDE STABILITY ----
df_pep = pd.read_csv(PEP_FILE)
freq_pct = dict(zip(df_pep["peptide"], df_pep["selection_freq_pct"]))
prot_map = dict(zip(df_pep["peptide"], df_pep["protein"]))

# ---- FIGURE ----
fig = plt.figure(figsize=(14,6), dpi=150)
gs = GridSpec(1,2,figure=fig, wspace=0.35)

# Panel A
ax1 = fig.add_subplot(gs[0,0])
ax1.plot(fpr, tpr, lw=2.5, color="#2C3E50",
         label=f"AUC = {final_auc:.3f}")
ax1.plot([0,1],[0,1],"--",color="gray")
ax1.scatter([1-spec],[sens],color="red",s=60,
            label=f"Sens={sens:.2f}, Spec={spec:.2f}")
ax1.set_xlabel("1 - Specificity")
ax1.set_ylabel("Sensitivity")
ax1.set_title("A", loc="left", fontweight="bold")
ax1.legend()
ax1.grid(alpha=0.3)

# ============================================================
# PANEL B — Stability Frequency (≥20% only)
# ============================================================

ax2 = fig.add_subplot(gs[0,1])

# Filter peptides with >=20%
filtered_items = [(pep,f) for pep,f in freq_pct.items() if f >= 20]

# Sort descending
filtered_items = sorted(filtered_items, key=lambda x:-x[1])

peps  = [x[0] for x in filtered_items]
freqs = [x[1] for x in filtered_items]

colors=[]
for pep,f in filtered_items:
    if f>=60:
        colors.append("#8E44AD" if prot_map[pep]=="gp41 env" else "#2980B9")
    elif f>=40:
        colors.append("#85929E")
    else:
        colors.append("#D5D8DC")  # ~20% bars

bars = ax2.bar(range(len(peps)), freqs, color=colors)

# ─────────────────────────────────────────────
# GROUPED LABELS (one per height level)
# ─────────────────────────────────────────────

unique_freqs = sorted(set(freqs), reverse=True)

for uf in unique_freqs:
    indices = [i for i, f in enumerate(freqs) if f == uf]
    if not indices:
        continue
    
    center = np.mean(indices)

    ax2.text(
        center,
        uf + 2,
        f"{int(uf)}%",
        ha="center",
        va="bottom",
        fontsize=9,
        fontweight="bold"
    )

# Threshold line
ax2.axhline(60,color="#E67E22",linestyle="--",linewidth=1.5)

ax2.set_ylabel("Selection Frequency (%)", fontweight="bold")
ax2.set_ylim(0,105)
ax2.set_xticks(range(len(peps)))
ax2.set_xticklabels(peps,rotation=60,ha="right",fontsize=7)

ax2.set_title("B", loc="left", fontweight="bold")
ax2.grid(axis="y",alpha=0.3)
ax2.set_title("B", loc="left", fontweight="bold")
ax2.grid(axis="y",alpha=0.3)

# ─────────────────────────────────────────────
# MAKE ALL TEXT BOLD
# ─────────────────────────────────────────────

for ax in [ax1, ax2]:

    ax.xaxis.label.set_fontweight("bold")
    ax.yaxis.label.set_fontweight("bold")

    for tick in ax.get_xticklabels():
        tick.set_fontweight("bold")
    for tick in ax.get_yticklabels():
        tick.set_fontweight("bold")

# Ensure peptide labels bold
for tick in ax2.get_xticklabels():
    tick.set_fontweight("bold")
plt.subplots_adjust(left=0.08, right=0.98, top=0.92, bottom=0.25)
plt.savefig(os.path.join(OUTPUT_PATH, "HIV_Figure_AB_20percent.png"),
            dpi=300, bbox_inches="tight")
plt.show()

# %% [code]
# SUPPLEMENTARY FIGURE S1 — RFECV Curve
# ═══════════════════════════════════════════════

OUT_S1 = os.path.join(OUTPUT_PATH, "Supplementary_Fig_S1_RFECV.png")

fig_s1 = plt.figure(figsize=(8,6), dpi=300)
ax = fig_s1.add_subplot(111)

nf_ = rfecv.cv_results_["n_features"]
ms_ = rfecv.cv_results_["mean_test_score"]
ss_ = rfecv.cv_results_["std_test_score"]

ax.plot(nf_, ms_, color="#2C3E50", lw=2.5)
ax.fill_between(nf_, ms_-ss_, ms_+ss_, alpha=0.2, color="#2C3E50")

ax.axvline(rfecv_n, color="#E67E22", lw=2, ls="--",
           label=f"Optimal features = {rfecv_n}")

ax.set_xlabel("Number of peptides", fontweight="bold")
ax.set_ylabel("Cross-validated AUC", fontweight="bold")

# 🔥 MAKE TICK NUMBERS BOLD
for tick in ax.get_xticklabels():
    tick.set_fontweight("bold")
for tick in ax.get_yticklabels():
    tick.set_fontweight("bold")

ax.legend()
ax.grid(alpha=0.3)

plt.tight_layout()

# Save
fig_s1.savefig(OUT_S1, bbox_inches="tight")

# 🔥 SHOW IN JUPYTER
plt.show()

# Optional: close AFTER showing
plt.close(fig_s1)

print("Supplementary Fig. S1 saved and displayed.")


# %% [code]
# ═══════════════════════════════════════════════
# SUPPLEMENTARY FIGURE S2 — AUC vs k (1% Rule)
# ═══════════════════════════════════════════════

OUT_S2 = os.path.join(OUTPUT_PATH, "Supplementary_Fig_S2_AUC_vs_k.png")

fig_s2 = plt.figure(figsize=(8,6), dpi=300)
ax = fig_s2.add_subplot(111)

ks   = K_VALUES
aucs = [mean_inner_auc[k] for k in ks]
stds = [std_inner_auc[k]  for k in ks]

# Plot curve
ax.plot(ks, aucs, "o-", color="#2C3E50", lw=2.5)
ax.fill_between(ks,
                [a-s for a,s in zip(aucs,stds)],
                [a+s for a,s in zip(aucs,stds)],
                alpha=0.2, color="#2C3E50",
                label="Mean ±1 SD")

# 1% equivalence band
max_auc = max(aucs)
tol = 0.01
lower_bound = max_auc - tol

ax.axhspan(lower_bound, max_auc,
           alpha=0.15, color="#27AE60",
           label="Within 1% of maximum AUC")

# Highlight selected k
ax.axvline(best_k, color="#E67E22", lw=2, ls="--",
           label=f"Selected k = {best_k}")
ax.scatter([best_k],
           [mean_inner_auc[best_k]],
           s=120, color="#E67E22", zorder=5)

# Labels
ax.set_xlabel("Number of peptides (k)", fontweight="bold")
ax.set_ylabel("Mean inner-CV AUC", fontweight="bold")
#ax.set_title("Supplementary Fig. S2 — Model Performance vs Feature Number",
#             fontweight="bold")

# 🔥 MAKE TICK NUMBERS BOLD
for tick in ax.get_xticklabels():
    tick.set_fontweight("bold")
for tick in ax.get_yticklabels():
    tick.set_fontweight("bold")
ax.set_ylim(0.85, 1.02)
ax.legend()
ax.grid(alpha=0.3)
plt.tight_layout()

fig_s2.savefig(OUT_S2, bbox_inches="tight")
# 🔥 SHOW IN JUPYTER
plt.show()

plt.close(fig_s2)

print("Supplementary Fig. S2 saved.")

# %% [code]


# %% [code]
```
