## Supplementary material for "Dual-Layer Serological Encoding: Integrating Pathogen-Derived and Host-Reactive Peptide Signatures Reveals Complementary Dimensions of HIV Immune Response": Code and Data: RRL_RF.ipynb

```
# %% [code]
"""
=============================================================================
 NESTED CROSS-VALIDATION — RANDOM FOREST + FEATURE REDUCTION
 RRL Peptide Library  |  n=105 samples
=============================================================================
 Sources:
   results_RRL_final.pkl
   df_samples_all.xls
   ListeSequenses_RRL_TBC_Test.xlsx

 High-dimensional configuration (p ≈ 2496, n=105)
=============================================================================
"""

import pickle, warnings, os
import numpy  as np
import pandas as pd
import matplotlib.pyplot as plt
import matplotlib.patches as mpatches
from matplotlib.gridspec import GridSpec
import seaborn as sns
from collections import Counter

from sklearn.ensemble import RandomForestClassifier
from sklearn.preprocessing import StandardScaler
from sklearn.model_selection import StratifiedKFold, permutation_test_score
from sklearn.metrics import (roc_auc_score, roc_curve,
                             confusion_matrix,
                             average_precision_score,
                             precision_recall_curve)
from sklearn.feature_selection import RFECV

# ─────────────────────────────────────────────────────────────
# SETTINGS
# ─────────────────────────────────────────────────────────────
import warnings
from sklearn.exceptions import (
    UndefinedMetricWarning,
    FitFailedWarning,
    ConvergenceWarning
)

warnings.filterwarnings("ignore", category=UndefinedMetricWarning)
warnings.filterwarnings("ignore", category=FitFailedWarning)
warnings.filterwarnings("ignore", category=ConvergenceWarning)
warnings.filterwarnings("ignore", message="X does not have valid feature names")
warnings.filterwarnings("ignore", message="Tight layout not applied")

# ⭐ THIS is the missing line for your endless warning:
warnings.filterwarnings(
    "ignore",
    message=".*sklearn.utils.parallel.delayed.*"
)
np.random.seed(42)
RS = 42
stable_thr = 40  # RRL more permissive (instead of 60)

# ─────────────────────────────────────────────────────────────
# PATHS
# ─────────────────────────────────────────────────────────────

DATA_PATH  = r"D:\Users\ha6057\Desktop\HIV_AI\Files for Analysis with Claud"
OUTPUT_PATH = DATA_PATH

PKL_FILE   = os.path.join(DATA_PATH, "results_RRL_final.pkl")
LABEL_FILE = os.path.join(DATA_PATH, "df_samples_all.xls")
SEQ_FILE   = os.path.join(DATA_PATH, "ListeSequenses_RRL_TBC_Test.xlsx")

OUT_MAIN_FIG    = os.path.join(OUTPUT_PATH, "RRL_RF_n105_Main.png")
OUT_STABLE_FIG  = os.path.join(OUTPUT_PATH, "RRL_RF_n105_StablePanel.png")
OUT_PEP_CSV     = os.path.join(OUTPUT_PATH, "RRL_RF_n105_peptides_ranked.csv")
OUT_OOF_CSV     = os.path.join(OUTPUT_PATH, "RRL_RF_n105_oof_predictions.csv")

# ─────────────────────────────────────────────────────────────
# 1) LOAD DATA
# ─────────────────────────────────────────────────────────────

print("\nLoading RRL dataset...")

with open(PKL_FILE, "rb") as f:
    rrl_data = pickle.load(f)

df_meta = pd.read_csv(LABEL_FILE)

label_map = (
    df_meta[df_meta["HIV_status"].isin(["HIV+","HIV-"])]
    .set_index("ID")["HIV_status"].to_dict()
)

overlap = sorted(set(rrl_data.keys()) & set(label_map.keys()))

rows = []
for sid in overlap:
    d = rrl_data[sid]
    row = dict(zip(d["ID"], d["MedianValue"]))
    row["SampleID"] = sid
    row["label"] = label_map[sid]
    rows.append(row)

feat_df = pd.DataFrame(rows)

META = ["SampleID","label"]
pep_cols = [c for c in feat_df.columns if c not in META]

X = feat_df[pep_cols].values.astype(np.float64)
y = (feat_df["label"] == "HIV+").astype(int).values

n, p = X.shape

print(f"Samples: {n}")
print(f"Peptides: {p}")

# ─────────────────────────────────────────────────────────────
# 2) ADAPTIVE K GRID (log scale)
# ─────────────────────────────────────────────────────────────

max_k = min(p, 1200)

K_VALUES = np.unique(
    np.round(np.logspace(np.log10(5), np.log10(max_k), 14)).astype(int)
).tolist()

print("k-grid:", K_VALUES)

# ─────────────────────────────────────────────────────────────
# 3) NESTED CV
# ─────────────────────────────────────────────────────────────

outer_cv = StratifiedKFold(n_splits=5, shuffle=True, random_state=RS)
inner_cv = StratifiedKFold(n_splits=5, shuffle=True, random_state=RS)

outer_probs = np.zeros(n)
fold_pep_idx = []
fold_importances = []

for fold_i, (tr_idx, te_idx) in enumerate(outer_cv.split(X, y)):

    X_tr, X_te = X[tr_idx], X[te_idx]
    y_tr, y_te = y[tr_idx], y[te_idx]

    sc = StandardScaler()
    Xtr_sc = sc.fit_transform(X_tr)
    Xte_sc = sc.transform(X_te)

    rf_rank = RandomForestClassifier(
        n_estimators=600,
        class_weight="balanced",
        max_features=0.2,
        min_samples_leaf=3,
        random_state=RS,
        n_jobs=1
    )

    rf_rank.fit(Xtr_sc, y_tr)

    imp = rf_rank.feature_importances_
    rank = np.argsort(-imp)

    fold_importances.append(imp.copy())

    k_auc = {}

    for k in K_VALUES:

        Xtr_k = Xtr_sc[:, rank[:k]]
        scores = []

        for in_tr, in_val in inner_cv.split(Xtr_k, y_tr):

            if len(np.unique(y_tr[in_val])) < 2:
                continue

            rf_in = RandomForestClassifier(
                n_estimators=400,
                class_weight="balanced",
                max_features=0.2,
                min_samples_leaf=3,
                random_state=RS,
                n_jobs=1
            )

            rf_in.fit(Xtr_k[in_tr], y_tr[in_tr])
            pv = rf_in.predict_proba(Xtr_k[in_val])[:,1]
            scores.append(roc_auc_score(y_tr[in_val], pv))

        k_auc[k] = np.mean(scores) if scores else 0

    best_v = max(k_auc.values())
    k_opt = min([k for k,v in k_auc.items() if v >= best_v - 0.01])

    fold_pep_idx.append(rank[:k_opt].tolist())

    rf_final = RandomForestClassifier(
        n_estimators=800,
        class_weight="balanced",
        max_features=0.2,
        min_samples_leaf=3,
        random_state=RS,
        n_jobs=1
    )

    rf_final.fit(Xtr_sc[:, rank[:k_opt]], y_tr)
    outer_probs[te_idx] = rf_final.predict_proba(Xte_sc[:, rank[:k_opt]])[:,1]

    print(f"Fold {fold_i+1} | k_opt={k_opt}")

# ─────────────────────────────────────────────────────────────
# 4) PERFORMANCE
# ─────────────────────────────────────────────────────────────

final_auc = roc_auc_score(y, outer_probs)
print("\nFinal AUC:", round(final_auc,4))

# ─────────────────────────────────────────────────────────────
# 5) STABILITY
# ─────────────────────────────────────────────────────────────

freq_cnt = Counter()
for idx_l in fold_pep_idx:
    freq_cnt.update(idx_l)

stable_idx = [i for i,c in freq_cnt.items()
              if (100*c/5) >= stable_thr]

stable_peps = [pep_cols[i] for i in stable_idx]

print("Stable peptides:", len(stable_peps))

# ─────────────────────────────────────────────────────────────
# 6) RFECV (safe for p=2496)
# ─────────────────────────────────────────────────────────────

sc_rfecv = StandardScaler()
X_sc = sc_rfecv.fit_transform(X)

rfecv = RFECV(
    RandomForestClassifier(
        n_estimators=400,
        class_weight="balanced",
        max_features=0.2,
        min_samples_leaf=3,
        random_state=RS,
        n_jobs=1
    ),
    step=50,
    cv=StratifiedKFold(5, shuffle=True, random_state=RS),
    scoring="roc_auc",
    min_features_to_select=10
)

rfecv.fit(X_sc, y)

print("RFECV optimal features:", rfecv.n_features_)

# ─────────────────────────────────────────────────────────────
# 7) SAVE OOF PREDICTIONS
# ─────────────────────────────────────────────────────────────

pd.DataFrame({
    "sample": feat_df["SampleID"],
    "true_label": y,
    "oof_prob": outer_probs
}).to_csv(OUT_OOF_CSV, index=False)

print("OOF saved.")
print("\nRRL analysis complete.")

import winsound
winsound.Beep(390, 700)

# %% [code]
# ─────────────────────────────────────────────────────────────
# FIGURE — RRL RESULTS (Styled Like Pathogen Figure)
# ─────────────────────────────────────────────────────────────

import matplotlib.pyplot as plt
from matplotlib.gridspec import GridSpec
from sklearn.metrics import roc_curve, confusion_matrix
import numpy as np
import os

# ─────────────────────────────────────────────
# COLORS (MATCH PREVIOUS FIGURE STYLE)
# ─────────────────────────────────────────────
C = dict(
    main="#2C3E50",      # ROC curve
    pos="#E74C3C",       # red operating point
    acc="#E67E22",       # threshold line
    purple="#8E44AD",    # >=60%
    grey="#7F8C8D",      # 40%
    light="#BDC3C7"      # 20%
)

fig = plt.figure(figsize=(14,6), dpi=150)
gs = GridSpec(1,2,figure=fig, wspace=0.35)

# ============================================================
# PANEL A — ROC
# ============================================================
ax1 = fig.add_subplot(gs[0,0])

fpr, tpr, thresholds = roc_curve(y, outer_probs)

# Youden index
j = np.argmax(tpr - fpr)
opt_thr = thresholds[j]

y_pred = (outer_probs >= opt_thr).astype(int)
tn, fp, fn, tp = confusion_matrix(y, y_pred).ravel()

sens = tp / (tp + fn)
spec = tn / (tn + fp)

# ROC
ax1.plot(fpr, tpr, lw=3, color=C["main"],
         label=f"AUC = {final_auc:.3f}")

# Diagonal
ax1.plot([0,1],[0,1],"--",color="gray", linewidth=1.5)

# Operating point
ax1.scatter([1-spec], [sens],
            color=C["pos"], s=100, zorder=5,
            label=f"Sens={sens:.2f}, Spec={spec:.2f}")

ax1.set_xlabel("1 - Specificity", fontweight="bold")
ax1.set_ylabel("Sensitivity", fontweight="bold")
ax1.set_title("A", loc="left", fontweight="bold", fontsize=14)

ax1.legend(frameon=False)
ax1.grid(alpha=0.25)

# ============================================================
# PANEL B — Stability Frequency (Grouped Labels)
# ============================================================

ax2 = fig.add_subplot(gs[0,1])

freq_pct = {pep_cols[i]: 100*freq_cnt[i]/5
            for i in freq_cnt}

sorted_items = sorted(freq_pct.items(),
                      key=lambda x:-x[1])

peps  = [x[0] for x in sorted_items]
freqs = [x[1] for x in sorted_items]

# Color logic
colors = []
for f in freqs:
    if f >= 60:
        colors.append(C["purple"])
    elif f >= 40:
        colors.append(C["grey"])
    else:
        colors.append(C["light"])

bars = ax2.bar(range(len(peps)), freqs, color=colors)

# Threshold line
stable_thr = 40
ax2.axhline(stable_thr,
            color=C["acc"],
            linestyle="--",
            linewidth=2)

# ─────────────────────────────────────────────
# GROUPED LABEL LOGIC
# ─────────────────────────────────────────────
unique_freqs = sorted(set(freqs), reverse=True)

for uf in unique_freqs:
    indices = [i for i, f in enumerate(freqs) if f == uf]
    if len(indices) == 0:
        continue
    
    # Center of the group
    center = np.mean(indices)
    
    ax2.text(
        center,
        uf + 1,
        f"{int(uf)}%",
        ha="center",
        va="bottom",
        fontsize=10,
        fontweight="bold"
    )

ax2.set_ylabel("Selection Frequency (%)", fontweight="bold")
ax2.set_ylim(0, 105)

ax2.set_xticks(range(len(peps)))
ax2.set_xticklabels(peps,
                    rotation=60,
                    ha="right",
                    fontsize=7,
                    fontweight="bold")

ax2.set_title("B", loc="left", fontweight="bold", fontsize=14)
ax2.grid(axis="y", alpha=0.25)

# Make tick labels bold
for ax in [ax1, ax2]:
    for tick in ax.get_xticklabels():
        tick.set_fontweight("bold")
    for tick in ax.get_yticklabels():
        tick.set_fontweight("bold")

# ─────────────────────────────────────────────
# SAVE
# ─────────────────────────────────────────────
fig.savefig(
    os.path.join(OUTPUT_PATH, "RRL_RF_n105_Styled.png"),
    dpi=300,
    bbox_inches="tight"
)

plt.show()

print("Styled figure saved.")

# %% [code]
import numpy as np
import matplotlib.pyplot as plt
from matplotlib.gridspec import GridSpec
from sklearn.metrics import roc_curve, confusion_matrix, roc_auc_score

# ------------------------------------------------------------
# COLOR PALETTE (same as HIV figure)
# ------------------------------------------------------------
C = dict(
    main="#2C3E50",
    pos="#E74C3C",
    purple="#8E44AD",
    darkgrey="#7F8C8D",
    lightgrey="#BDC3C7",
    acc="#E67E22"
)

# ------------------------------------------------------------
# CREATE FIGURE WITH MATCHING PROPORTIONS
# ------------------------------------------------------------
fig = plt.figure(figsize=(14,5))

gs = GridSpec(
    1, 2,
    figure=fig,
    width_ratios=[1, 1.4],
    wspace=0.25
)

ax1 = fig.add_subplot(gs[0,0])
ax2 = fig.add_subplot(gs[0,1])

# ============================================================
# PANEL A — ROC
# ============================================================

fpr, tpr, thresholds = roc_curve(y, outer_probs)
auc = roc_auc_score(y, outer_probs)

j = np.argmax(tpr - fpr)
opt_thr = thresholds[j]

y_pred = (outer_probs >= opt_thr).astype(int)
tn, fp, fn, tp = confusion_matrix(y, y_pred).ravel()

sens = tp / (tp + fn)
spec = tn / (tn + fp)

ax1.plot(fpr, tpr, lw=2.5, color=C["main"],
         label=f"AUC = {auc:.3f}")

ax1.plot([0,1],[0,1],"--",color="gray")

ax1.scatter([1-spec], [sens],
            color=C["pos"], s=120, zorder=5,
            label=f"Sens={sens:.2f}, Spec={spec:.2f}")

ax1.set_xlabel("1 - Specificity", fontweight="bold")
ax1.set_ylabel("Sensitivity", fontweight="bold")
ax1.set_title("A", loc="left", fontweight="bold", fontsize=16)

ax1.set_xlim(0,1)
ax1.set_ylim(0,1)

ax1.legend(frameon=False)
ax1.grid(alpha=0.25)

# ============================================================
# PANEL B — Stability Frequency
# ============================================================

freq_pct = {pep_cols[i]: 100*freq_cnt[i]/5
            for i in freq_cnt}

sorted_items = sorted(freq_pct.items(),
                      key=lambda x:-x[1])

peps  = [x[0] for x in sorted_items]
freqs = [x[1] for x in sorted_items]

colors = []
for f in freqs:
    if f >= 60:
        colors.append(C["purple"])
    elif f >= 40:
        colors.append(C["darkgrey"])
    else:
        colors.append(C["lightgrey"])

bars = ax2.bar(range(len(peps)), freqs, color=colors)

# Threshold line
ax2.axhline(40, color=C["acc"],
            linestyle="--", linewidth=2)

# Grouped labels
unique_freqs = sorted(set(freqs), reverse=True)

for uf in unique_freqs:
    idx = [i for i, f in enumerate(freqs) if f == uf]
    center = np.mean(idx)
    ax2.text(center, uf + 1.5,
             f"{int(uf)}%",
             ha="center",
             va="bottom",
             fontsize=12,
             fontweight="bold")

# ---------- FIX DUPLICATE TICKS ----------
ax2.set_ylim(0, 100)
ax2.set_yticks([0,20,40,60,80,100])
ax2.set_yticklabels([0,20,40,60,80,100], fontweight="bold")

ax2.set_ylabel("Selection Frequency (%)",
               fontweight="bold")

ax2.set_xticks(range(len(peps)))
ax2.set_xticklabels(peps,
                    rotation=60,
                    ha="right",
                    fontsize=7,
                    fontweight="bold")

ax2.set_title("B", loc="left",
              fontweight="bold",
              fontsize=16)

ax2.grid(axis="y", alpha=0.25)

# ------------------------------------------------------------
plt.tight_layout()
plt.show()

# %% [code]
# ─────────────────────────────────────────────────────────────
# HEATMAP — Stable Peptides (Z-scored) — Clean Journal Version
# ─────────────────────────────────────────────────────────────

import numpy as np
import pandas as pd
import matplotlib.pyplot as plt
import seaborn as sns
from matplotlib.colors import TwoSlopeNorm

# ---------- 1. Prepare matrix (stable peptides only)

stable_idx = [pep_cols.index(p) for p in stable_peps]
X_stable   = X[:, stable_idx]

# Z-score across samples (per peptide)
X_z = (X_stable - X_stable.mean(axis=0)) / X_stable.std(axis=0)

# Clip at ±3 SD
X_z = np.clip(X_z, -3, 3)

# ---------- 2. Order samples

df_heat = pd.DataFrame(X_z, columns=stable_peps)
df_heat["HIV"]  = y
df_heat["prob"] = outer_probs

df_pos = df_heat[df_heat["HIV"] == 1].sort_values("prob", ascending=False)
df_neg = df_heat[df_heat["HIV"] == 0].sort_values("prob", ascending=False)

df_sorted = pd.concat([df_pos, df_neg])
heat_matrix = df_sorted[stable_peps].T.values

n_pos = df_pos.shape[0]
n_neg = df_neg.shape[0]

# ---------- 3. Create heatmap

plt.figure(figsize=(16, 6), dpi=150)

norm = TwoSlopeNorm(vmin=-3, vcenter=0, vmax=3)

ax = sns.heatmap(
    heat_matrix,
    cmap="RdBu_r",
    norm=norm,
    xticklabels=False,
    yticklabels=stable_peps,
    cbar_kws={"label": "Z-score (IgG signal)"}
)

# ---------- 4. Vertical separator

plt.axvline(n_pos, color="black", linewidth=2)

# ---------- 5. Make all labels bold & black

# Y-axis label
ax.set_ylabel("Peptides", fontsize=12, fontweight="bold", color="black")

# Remove X-axis label
ax.set_xlabel("")

# Peptide labels
for label in ax.get_yticklabels():
    label.set_fontweight("bold")
    label.set_color("black")

# Remove X ticks completely
ax.set_xticks([])

# Colorbar formatting
cbar = ax.collections[0].colorbar
cbar.ax.yaxis.label.set_fontweight("bold")
cbar.ax.yaxis.label.set_color("black")

for t in cbar.ax.get_yticklabels():
    t.set_fontweight("bold")
    t.set_color("black")

# ---------- 6. Remove title (do nothing)

plt.tight_layout()

# ---------- 7. Save figure

plt.savefig(
    "HIV_StablePanel_Heatmap_Clean.png",
    dpi=300,
    bbox_inches="tight"
)

plt.show()

# %% [code]
print("\nFinal stable peptides:")
for p in stable_peps:
    print(p)

# %% [code]
print("RFECV optimal features:", rfecv.n_features_)

# %% [code]


# %% [code]


# %% [code]
```
